# Respiratory mucosal administration of DNA aptamer nanomaterials protects against antigenically diverse SARS-CoV-2 variants

**DOI:** 10.1101/2024.05.31.596896

**Authors:** Michael R. D’Agostino, Jiuxing Li, Zijie Zhang, Jimmy Gu, Art Marzok, Jann Ang, Katherine E. Bujold, Sam Afkhami, Xiaohu Xia, Yingfu Li, Matthew S. Miller

**Author notes:** Denotes co-corresponding authors. Denotes co-first authors (Michael R. D’Agostino, Jiuxing Li).

## Abstract

The ongoing COVID-19 pandemic has highlighted the need for innovative therapeutic strategies to combat rapidly evolving pathogens that challenge the efficacy of traditional vaccines and monoclonal antibody treatments. Here, we explored the potential of TMSA52, a previously described homotrimeric DNA aptamer as a universal prophylactic and therapeutic agent against SARS-CoV-2. TMSA52 demonstrates exceptional binding affinities and broad neutralization against diverse SARS-CoV-2 variant spike proteins that are further enhanced through multimerization onto lamellar iridium nanoplates. Respiratory mucosal delivery of TMSA52 nanomaterials was well-tolerated. Surprisingly, TMSA52 offered potent protection from infection with ancestral SARS-CoV-2 on-par with monoclonal antibodies, and superior protection against antigenically distant SARS-CoV-2 variants. These findings establish DNA aptamers as a promising, cost-effective, and scalable alternative to traditional monoclonal antibody therapies. This study underscores the potential of aptamer-based platforms as a next-generation strategy to enhance global pandemic preparedness and expand our arsenal of infectious disease countermeasures.

## INTRODUCTION

Since the emergence of the COVID-19 pandemic, SARS-CoV-2 has claimed over seven million lives and continues to adversely affect health systems across the globe (*1*). Although first-generation and updated booster vaccines have been critical in mitigating the impact of COVID-19, their efficacy is jeopardized by both the continued antigenic evolution of SARS-CoV-2 and waning of vaccine-induced immunity (*2*). This has resulted in ongoing hospitalizations and deaths associated with COVID-19, which remains particularly evident in immunocompromised and elderly populations, who often generate suboptimal immunity following vaccination (*3*).

Monoclonal antibodies (mAb) specific for the spike protein of SARS-CoV-2 were among the few therapies authorized to treat COVID-19, with numerous studies confirming their efficacy in reducing hospitalizations, severe disease, and fatalities (*4, 5*). However, many of these antibodies rapidly lost efficacy due to the emergence of new variants of concern (VoC) such as XBB.1.5 and JN.1, with recent findings indicating that Sotrovimab, one of the last effective mAbs, can no longer neutralize the predominantly circulating variants (*2, 6*). This has necessitated development of new mAbs that have specificity for the newest variants of SARS-CoV-2. The limitations of mAb therapies for COVID-19, including their complex and relatively expensive manufacturing processes and the need for intravenous (i.v.) administration, present significant challenges, resulting in high cost of treatment and limited accessibility (*7, 8*). Likewise, there is an urgent need for effective prophylactic options capable of protecting high-risk individuals who respond poorly to vaccines (e.g., immunocompromised, elderly, etc.). This highlights the need for next-generation COVID-19 prophylactics and therapeutics that are effective against both current and future SARS-CoV-2 VoC that have cheaper and more efficient manufacturing processes, and that harness more practical delivery methods to enhance efficacy, cost-effectiveness, and accessibility. As the respiratory tract is the primary site of SARS-CoV-2 infection, it is pragmatic to design new treatments for intranasal/inhaled aerosol delivery as this method of delivery targets therapies directly to the initial site of infection, the respiratory mucosa (RM).

DNA aptamers are single-stranded oligonucleotides derived from complex random pools by systematic evolution of ligands by exponential enrichment (SELEX) for their ability to bind molecules-of-interest such as proteins (*9*). In contrast to antibodies, DNA aptamers possess superior stability and can be commercially scaled for a fraction of the cost (*10*). DNA aptamers have great utility in the field of molecular diagnostics, and when designed to target the receptor-binding sites utilized by pathogens for host entry, may offer promising prophylactic/therapeutic activity (*11–13*). Our group has previously described a homotrimeric DNA aptamer, TMSA52, composed of three single-stranded DNA molecules trimerized around a phosphoramidite trebler that possesses binding affinities ranging from 8.8 pM for the ancestral SARS-CoV-2 spike protein, to 23.7 pM for the Omicron BA.1 spike protein (*14, 15*). Its ability to bind antigenically diverse spike proteins from a wide range of SARS-CoV-2 VoC with exquisite sensitivity and specificity suggested that TMSA52 might have therapeutic applications in addition to its utility as a molecular diagnostic tool (*15*).

In this study, we tested whether TMSA52 could be used as a universal agent to protect against SARS-CoV-2. Herein, we report on novel topical administration directly to the respiratory mucosa of a DNA aptamer as an anti-infective prophylactic agent against SARS-CoV-2. We report that the trimerization of MSA52, and further multimerization of TMSA52 onto lamellar iridium nanoplates (IrNP@TMSA52) increases the stability of the aptamer, and further enhances its ability to neutralize SARS-CoV-2 XBB.1.5. When administered intranasally (i.n.) to mice, TMSA52 and IrNP@TMSA52 offered complete protection against ancestral SARS-CoV-2 and contemporary VoC infection, preventing morbidity, mortality, and lung pathology, while also reducing lung viral burden. Importantly, TMSA52 does not induce an overt inflammatory response upon pulmonary administration before or after SARS-CoV-2 infection, offering a tolerable safety profile required for inhaled therapeutics. The development of novel, broadly protective aptamer nanomaterials against viral pathogens expands the arsenal of antiviral countermeasures beyond vaccines and monoclonal antibodies and highlights the advantages of aptamer-based biologics as a prospective mucosal treatment against SARS-CoV-2 due to their customizable synthesis, ambient temperature stability, and amenability for topical administration.

## RESULTS

### Multimerization of the universal SARS-CoV-2 aptamer MSA52 enhances neutralization of SARS-CoV-2

All SARS-CoV-2 mAbs were designed to bind to the SARS-CoV-2 spike and consequently neutralize virus. To date, the majority of these mAb have steadily lost their efficacy, owing to the continued antigenic drift of SARS-CoV-2. Sotrovimab remained one of the last mAb therapies capable of efficiently neutralizing Omicron BA.1 (*16*). Despite its durable efficacy against a wide array of pre-Omicron SARS-CoV-2 variants, the continued evolution of SARS-CoV-2 has concomitantly eroded the efficacy of Sotrovimab. Sotrovimab was unable to neutralize XBB.1.5 even at the highest FRNT_50_ value tested of ≥ 50,000 ng/mL, and the contemporary JN.1 variant demonstrated near complete loss in neutralization, potentially due to a D339H mutation in the spike protein (*17, 18*).

To date, several multimeric aptamers based on DNA nanostructures, DNA origami, or gold nanoparticles have been reported that neutralize ancestral SARS-CoV-2. However, none of these multimeric aptamers can neutralize SARS-CoV-2 variants without a substantial drop in efficacy (*11, 19, 20*). As an example, a spherical cocktail neutralizing aptamer-gold nanoparticle (SNAP) was capable of neutralizing pseudoviruses with spike proteins derived from the ancestral SARS-CoV-2, and variants harboring a combination of N501Y, D614G, K417N, and E484K mutations. The jump in IC_50_ values between the ancestral SARS-CoV-2 and these early variant spike proteins was as substantial as 432-fold (*21*). Our group has previously described that trimerization of MSA52 to TMSA52 increases its affinity for spike, and that TMSA52 binds pseudoviruses with the spike protein of eight SARS-CoV-2 variants, experiencing no more than a 4.3-fold loss in affinity (*15*).

Given that trimerizing MSA52 into TMSA52 using a phosphoramidite trebler increases its affinity for the spike protein of numerous SARS-CoV-2 VoC, we first sought to assess if TMSA52 retained increased neutralization against SARS-CoV-2 BA.5 variant spike protein-expressing pseudoviruses. To this end, serial dilutions of MSA52 or TMSA52 were incubated with an enhanced green fluorescent protein (eGFP)-expressing pseudovirus expressing the SARS-CoV-2 Omicron BA.5 spike. Angiotensin-converting enzyme 2 (ACE2) expressing HEK 293T cells were infected with the BA.5 spike-expressing pseudoviruses following co-incubation with various concentrations of aptamer nanomaterials and after 48 hours the ratio of infected GFP-positive cells to total cells was quantified (Figure S1A, B). The IC_50_ was calculated by plotting the ratio of infected cells against the concentration of aptamer nanomaterials. In comparison to the monomeric aptamer MSA52, trimeric TMSA52 exhibited a 52-fold decrease in IC_50_ (Figure 1A). Given that trimerization increased the neutralization potency of MSA52, we next sought to further increase the valency of our aptamer nanomaterials by conjugating them onto lamellar iridium nanoplates (IrNPs). The novel one-pot synthesis process is described herein, and the physical properties of the IrNPs were extensively characterized (Figure S2 and S3). To facilitate the conjugation of aptamers, IrNPs were coated with streptavidin by physical adsorption, followed by attachment of biotinylated trimeric aptamer (TMSA52-B) (Table S1) to obtain IrNP@TMSA52 (Figure 1B, Figure S4A). To quantify the number of TMSA52-B aptamers on each IrNP, a FAM-labelled antisense sequence (FAM-AS, Table S1) complementary to TMSA52-B was hybridized with IrNP@TMSA52 conjugates. After removing excess FAM-AS, IrNP@TMSA52 conjugates were denatured at 90°C in PBS containing 8 M urea to release FAM-AS. The fluorescent intensity of the supernatant was measured to quantify the amount of TMSA52-B bound to each IrNP using a standard curve. Saturation of the surface of each IrNP with TMSA52-B was achieved at 5.17 × 10^5^ aptamer copies (Figure S4B). Notably, affixing TMSA52 onto the IrNPs markedly increased aptamer stability in the presence of serum nucleases, likely due to increased steric hindrance (Figure S4C, D). IrNP@TMSA52 retained its ability to bind to- and neutralize pseudoviruses expressing a range of SARS-CoV-2 variant spike proteins (Figure S4E, F). As a comparator, a gold nanoparticle AuNP conjugated aptamer (AuNP@TMSA52) was utilized as a relevant nanoparticle control, owing to the widespread usage of gold nanoparticles clinically. The conjugation of MSA52 onto IrNPs and AuNPs resulted in IC_50_ reductions of 92.7-fold and 18.9-fold respectively (Figure 1C, Figure S1C, D), likely due to increased stability and improved avidity arising from multivalent interactions with the spike protein. Moreover, the conjugation of TMSA52 on IrNPs and AuNPs resulted in a further 16.6-fold and 2.7-fold decrease in IC_50_, respectively (Figure 1C, Figure S1E, F). IrNPs alone had not appreciable neutralization activity against BA.5 pseudovirus (Figure S1G). The enhanced neutralization of aptamer-conjugated IrNPs relative to AuNPs for both MSA52 and TMSA52 may be due to the increased steric hindrance of receptor binding provided by the large, 2-dimensional structure of IrNPs. Collectively, these data reinforce prior evidence that trimeric TMSA52 outperforms monomeric MAS52, and furthermore suggest that IrNP-based nanomaterials are superior to AuNP-based nanomaterials at neutralizing SARS-CoV-2 BA.5 spike pseudoviruses.

**Fig. 1.**
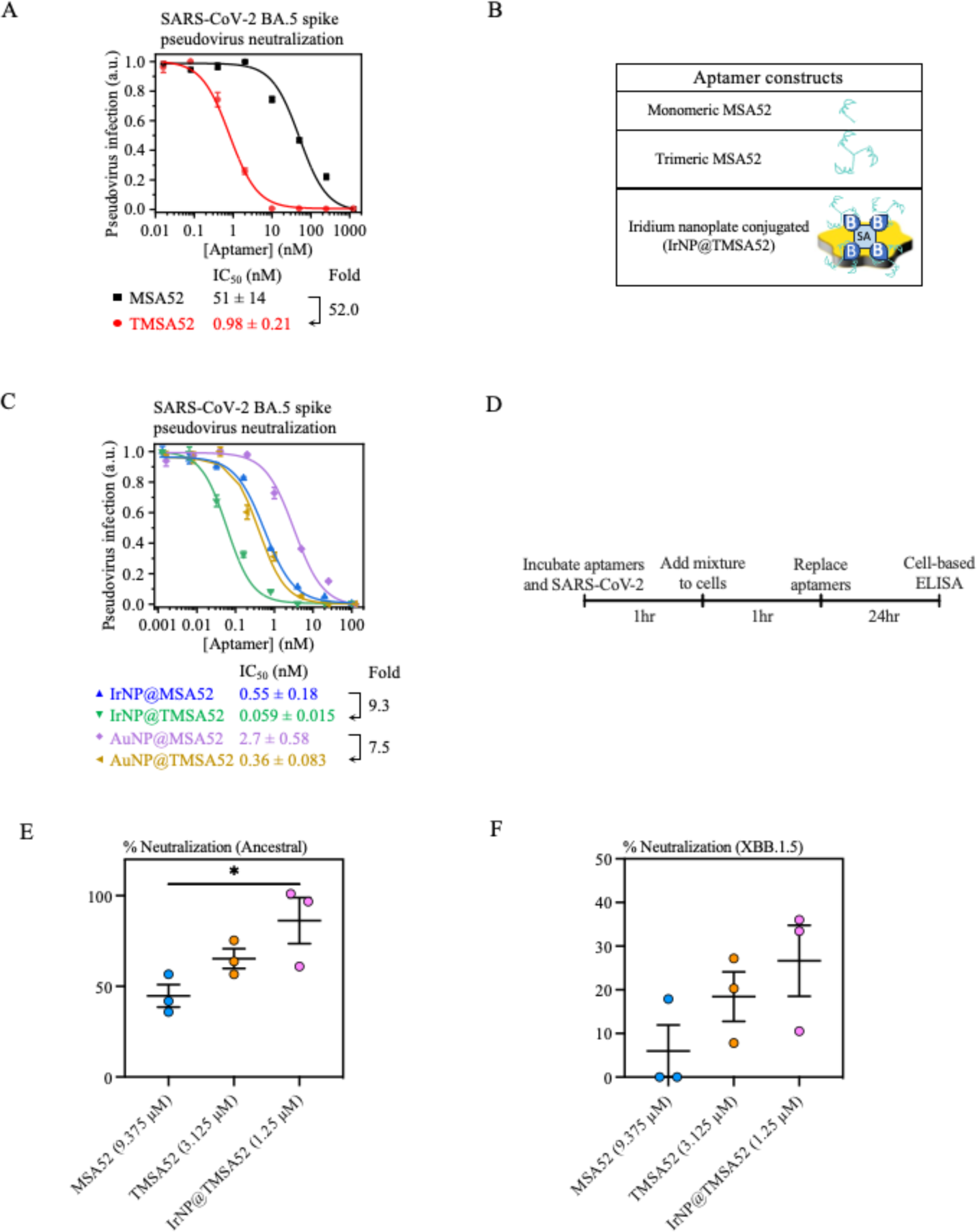
*In vitro* neutralization of SARS-CoV-2 variants by aptamers and aptamer-conjugated nanomaterials. (**A**) Determination of IC_50_ values for different monomeric MSA52 and TMSA52 for the neutralization of BA.5 spike expressing pseudovirus. (**B**) Schema of monomeric MSA52, trimeric TMSA52, and biotinylated TMSA52 conjugated on streptavidin coated IrNPs by biotin (B) - streptavidin (SA) interaction. (**C**) Determination of IC_50_ values for different aptamer-conjugated nanomaterials for the neutralization of BA.5 spike expressing pseudovirus. (**D**) Experimental schema for SARS-CoV-2 neutralization assay. (**E**) Neutralization plots against ancestral SARS-CoV-2 using 9.375 µM MSA52, 3.125µM TMSA52, or 1.25µM IrNP@TMSA52. (**F**) Neutralization plots against SARS-CoV-2 Omicron XBB.1.5 using 9.375 µM MSA52, 3.125µM TMSA52, or 1.25µM IrNP@TMSA52. Statistical analysis for panel E was a one-way ANOVA with Tukey’s multiple comparisons test (*P < 0.05*). Data is representative of one independent experiment (**A, C**) or pooled from three independent experiments (**E, F**).

We have thus far confirmed that IrNP@TMSA52 outperforms alternate aptamer nanomaterials at neutralizing pseudoviruses expressing the spike proteins from an array of SARS-CoV-2 variants. We next sought to assess the potency of TMSA52 in neutralizing *bona fide* SARS-CoV-2 by utilizing a well-established *in vitro* neutralization assay (Figure 1D). In congruence with the pseudovirus neutralization assay, TMSA52 demonstrated superior neutralization to MSA52, and IrNP@TMSA52 further increased neutralization against an isolate of the ancestral SARS-CoV-2 virus (Figure 1E). When tested against the contemporary XBB.1.5 variant MSA52 demonstrated a small 3.1-fold drop in neutralization potency relative to ancestral virus, while TMSA52 had a 2.6-fold drop in efficacy. Similar to the relative resilience of IrNP@TMSA52 to antigenic changes observed in the pseudovirus neutralization assay, we only observed a 2.5-fold decrease in neutralization against the SARS-CoV-2 XBB.1.5 variant (Figure 1F). The above data indicate that multimerization of the aptamer nanomaterials increased their ability to neutralize SARS-CoV-2, with TMSA52 offering superior neutralization relative to MSA52, and IrNP@TMSA52 providing the most robust neutralization against SARS-CoV-2. Furthermore, multimerization increased the resilience of the aptamer nanomaterials to antigenic changes in the spike protein and enhanced their ability to neutralize the antigenically distant XBB.1.5 variant.

### Respiratory mucosal prophylaxis with TMSA52 provides robust protection against lethal SARS-CoV-2 infection

Thus far we have demonstrated that aptamer nanomaterials are capable of binding to- and universally neutralizing multiple SARS-CoV-2 VoCs *in vitro*. To determine the clinical potential of our TMSA52-based nanomaterials, we next assessed their ability to confer protection against SARS-CoV-2 *in vivo*. To this end, BALB/c mice were i.n. administered either TMSA52, or IrNP@TMSA52. As controls, a subset of animals received either nuclease-free phosphate buffered saline (PBS; vehicle), a nonsense DNA aptamer with equivalent nucleotide ratios as TMSA52 but with abolished binding to SARS-CoV-2 spike, or IrNP alone. As a positive control the neutralizing antibody S309 (the parental antibody of the clinically approved Sotrovimab) was used (Figure 2A) (*22*). Two hours post-administration, all animals were infected with a lethal dose of a mouse-adapted strain of ancestral SARS-CoV-2 (MA10) and monitored for weight loss (a measure of morbidity) and mortality (*23*). A subset of animals was sacrificed 4 days post-infection (dpi) for enumeration of lung viral titers and for histopathological analysis of the lung. Animals treated with either TMSA52, IrNP@TMSA52, or S309 were wholly protected from weight loss, with all animals surviving lethal infection (Figure 2B, C). In contrast, animals which received either PBS, IrNP, or the nonsense DNA aptamer rapidly lost weight, with some animals reaching humane endpoint by 4 days post-infection (Figure 2B, C).

**Fig. 2.**
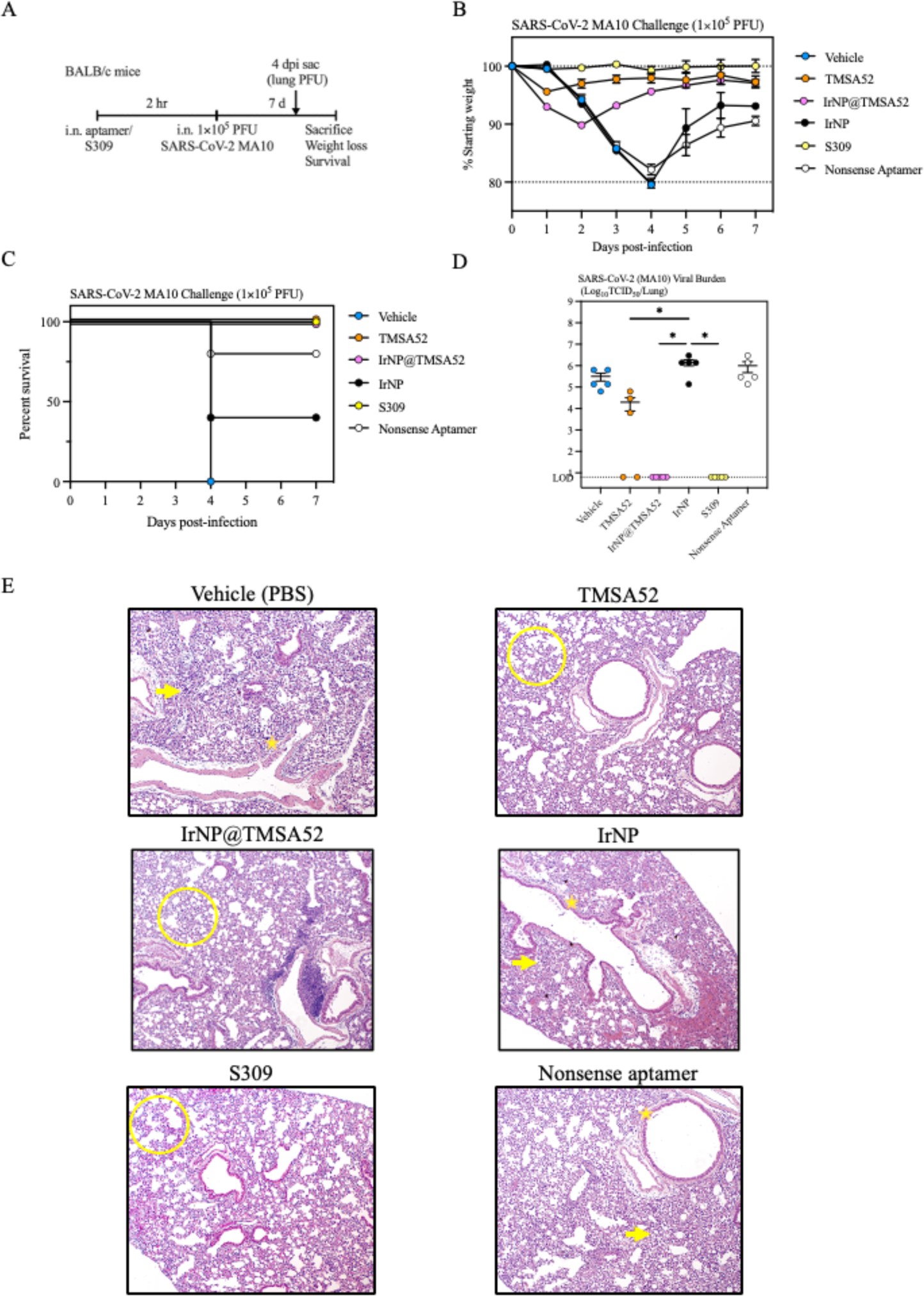
Protective assessment of intranasally delivered aptamers against lethal challenge with SARS-CoV-2. (**A**) Animals were treated intranasally (i.n.) with either nuclease-free PBS (vehicle), TMSA52 (SARS-CoV-2 aptamer), IrNP@TMSA52 (universal SARS-CoV-2 aptamer affixing on an iridium nanoplate), IrNP (iridium nanoplate alone), S309 (SARS-CoV-2 monoclonal antibody), or nonsense aptamer (a nonsense TMSA52 with equimolar A:T and C:G ratios with abrogated binding to SARS-CoV-2 spike) 2 hours prior to i.n. challenge with 1 × 10_5_ PFU of SARS-CoV-2 MA10. Animals were subsequently monitored for weight loss and mortality for seven days with a sub cohort sacrificed four days post-infection for lung viral titers and histopathology. (**B**) Weight loss following SARS-CoV-2 MA10 challenge. (**C**) Survival following SARS-CoV-2 MA10 challenge. (**D**) Lung viral burden of a cohort of animals (n = 5) sacrificed four days post-infection. (**E**) Hematoxylin and eosin-stained lung sections from a cohort of animals sacrificed four days post-infection with yellow arrows indicating areas of diffuse epithelial damage, yellow stars representing epithelial sloughing, and yellow circles representing healthy lung architecture. Data is presented as mean ± SEM. Statistical analysis for panel D was a one-way ANOVA with Tukey’s multiple comparisons test (*P < 0.05*). Data is representative of one independent experiment, n = 5 mice/group.

Within the subset of animals sacrificed at 4 dpi, all mice treated with S309 and IrNP@TMSA52 had no recoverable virus in the lungs. Two of five mice treated with TMSA52, had no recoverable viral titers from the lungs (Figure 2D). TMSA52 treated mice that did not clear the virus by 4 dpi had a 17-fold decrease in viral burden relative to vehicle-treated mice (1.9 × 10^4^ TCID_50_ vs. 3.2 × 10^5^ TCID_50_; Figure 2D). Mice treated with the nonsense aptamer or IrNP alone had comparable lung viral burden to vehicle-treated mice (Figure 2D). Importantly, animals treated with either the nonsense aptamer or IrNP alone has similar lung viral titers to PBS-treated control animals, indicating the protection afforded by TMSA52 and IrNP@TMSA52 required aptamer binding to SARS-CoV-2 virus.

Hematoxylin and eosin (H&E) staining was performed on lung sections from infected animals at 4 dpi, revealing prominent epithelial damage (yellow arrows) and epithelial sloughing (yellow stars) in the lungs of vehicle and IrNP treated mice (Figure 2E, top left, middle right). Mice treated with the nonsense aptamer exhibited similar pathological features, congruent with their substantial weight loss (Figure 2E, bottom right). As expected, S309 treated mice had preserved lung architecture (yellow circles) and showed minimal pulmonary damage (Figure 2E, bottom left). In alignment with the marked protection against weight loss and morbidity, lung histopathology revealed that TMSA52 and IrNP@TMSA52 treated animals showed drastically reduced epithelial damage and sloughing in comparison to vehicle, IrNP alone, and nonsense aptamer treated mice (Figure 2E, top right and middle left). Collectively, these data demonstrate that TMSA52, and IrNP@TMSA52 can protect mice against morbidity and mortality following lethal SARS-CoV-2 infection, comparable to a gold-standard monoclonal antibody treatment.

### Intranasal aptamer nanomaterial administration does not induce aberrant inflammatory responses before and after SARS-CoV-2 infection

Monoclonal antibodies are a safe and generally well-tolerated frontline therapy offered to individuals infected with SARS-CoV-2 that are at risk of progressing to severe disease. In contrast, extra-nuclear DNA is a canonical danger-associated molecular pattern (DAMP) that can announce the presence of cellular damage or a pathogen and can potently stimulate the immune system (*24*). The potential of causing a hyper-inflammatory response due to the presence of exogenous DNA is of importance in the lung, where a delicate balance must be struck to maintain the critical function of gas exchange.

To determine if respiratory mucosal aptamer nanomaterial administration induces an inflammatory response in the airways, BALB/c mice were treated i.n. with either nuclease-free PBS (vehicle), TMSA52, IrNP@TMSA52, or a nonsense aptamer that does not bind to the SARS-CoV-2 spike (Figure 3A). Two hours after aptamer nanomaterial treatment, one cohort of mice from each group were sacrificed prior to infection (0 hr), and bronchoalveolar lavage (BAL) was performed to assess airway cytokine responses, and the cellular compartment was subjected to comprehensive flow cytometric analysis to profile innate immune cell infiltration. A second cohort was infected with SARS-CoV-2 MA10 and sacrificed at 1 dpi (24 hr), and again BAL was performed to quantify airway cytokines and immune cell infiltrates (*25*). All vehicle treated mice had negligible levels of innate inflammatory cytokines at 2 hours post-treatment prior to infection (Figure 3B-G). Irrespective of treatment, no mice had increased levels of IL-1β, IFN-β1, or MCP-1 before infection (Figure 3B, E, G). Mice receiving TMSA52, had elevated IL-6, TNF-α, and MIP-1α in the airways, comparable to mice receiving nonsense aptamer prior to infection, but less than mice treated with IrNP@TMSA52 (Figure 3B-G). At 24 hours post-infection vehicle mice exhibited a modest increase in IL-6 and MIP-1α, comparable to mice receiving nonsense aptamer (Figure 3C, F). Mice receiving TMSA52 or IrNP@TMSA52 had increased levels of IL-1-β, IL-6, TNF-α, MIP-1-α and MCP-1 (Figures 3B, C, D, F, G). Notably, nonsense aptamer administration did not heighten the inflammatory response in the airways after infection to the same extent as TMSA52-based nanomaterials, indicating that the slight increase in inflammatory cytokines observed is dependent on the interaction between aptamer nanomaterial and the viral spike protein. Importantly, those responses correlate with protection, as TMSA52-treated groups all survived lethal challenge with minimal morbidity, while mice treated with nonsense aptamer experienced severe weight loss, with some mice reaching endpoint (Figure 2). No treatment group exhibited a consistent increase in IFN-β1 either before or after SARS-CoV-2 infection, indicating that respiratory mucosal administration of these aptamer-based nanomaterials did not induce a profound type I IFN response (Figure 3E).

**Fig. 3.**
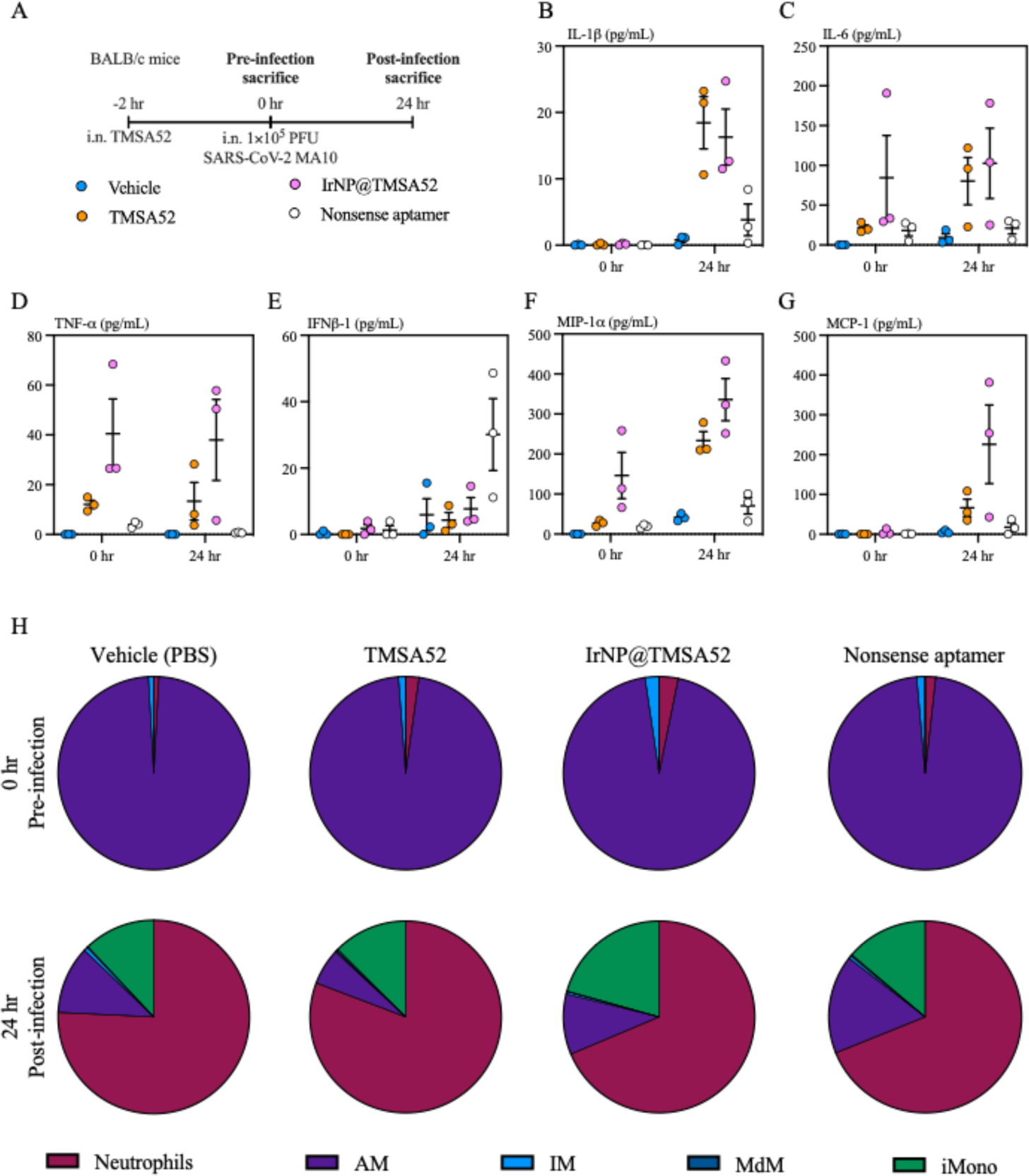
Airway inflammatory profiling following intranasal aptamer administration before and after SARS-CoV-2 challenge. (**A**) Animals were treated intranasally (i.n.) with either nuclease free-PBS (vehicle), TMSA52, IrNP@TMSA52, or a nonsense DNA aptamer. One cohort of mice was sacrificed two hours after treatment and bronchoalveolar lavage (BAL) was performed for immune profiling (0 hr pre-infection sacrifice). A second cohort from each treatment group was subsequently challenged with 1 × 10_5_ PFU of SARS-CoV-2 MA10 two hours after treatment and sacrificed 24 hours post-infection (24hr post-infection sacrifice) for airway cytokine multiplex analysis and flow cytometric analysis. (**C-E**) Dot-plots depicting bronchoalveolar lavage fluid cytokine concentrations of IL-1β (B), IL-6 (C), TNF-α (D), IFN-β1 (E), MIP-1α (F), and MCP-1 (G). (**H**) Pie charts depicting the average frequencies of live, CD45_+_ bronchoalveolar lavage neutrophils, alveolar macrophages (AM), interstitial macrophages (IM), monocyte-derived macrophages (MdM), and inflammatory monocytes before and after SARS-CoV-2 MA10 infection. Data is representative of one independent experiment, line and error bars represent mean ± SEM, pie charts represent the mean frequencies of Live, CD45_+_ BAL cells, n = 3 mice/group.

In addition to the cytokine milieu, BAL cells were subjected to flow cytometric analysis using a previously published panel (gating strategy Figure S5) (*25–28*). At 2 hours post-aptamer nanomaterial administration, but before infection (0 hr), 90-96 % of all live CD45^+^ cells in the airways of all mice regardless of treatment were alveolar macrophages (AM; Figure 3H, top). At 24 hours post-infection, all mice experienced a large influx of neutrophils, ranging from 63-73 % for nonsense aptamer treated mice to TMSA52 treated mice respectively (Figure 3H, bottom). In addition to neutrophils, following SARS-CoV-2 infection all treatment groups had elevated frequencies of Ly6C^+^ inflammatory monocytes, ranging from 11.1 % for vehicle treated mice to 17.8 % for IrNP@TMSA52 treated mice (Figure 3H, bottom). Taken together, these data indicate that respiratory mucosal administration with DNA aptamer nanomaterials induce minimal immune responses prior to infection, while elevated levels of IL-1β, IL-6, TNF-α, MIP-1α, and MCP-1 in TMSA52-treated groups post-infection correlates with protection.

### Protection offered by a single intranasal dose of TMSA52 persists for at least 24 hours

Existing RNA aptamer therapies are administered topically into the vitreous fluid of the eye, due to their transient half-life in the serum (*29*). The respiratory mucosa is bathed in cellular and acellular components which act to maintain the primary function of the lung – gas exchange. These barriers can often interfere with the delivery of therapeutics (*30*).

To determine if the protection offered by TMSA52 can persist for a protracted period in the airways, we administered TMSA52 i.n. to mice either 2 hours or 24 hours prior to SARS-CoV-2 MA10 challenge (Figure 4A). Additionally, we sought to assess whether lower doses of TMSA52 could provide commensurate protection to the standard dose of 258 µM used in earlier experiments. To this end, we included cohorts that received 25.8 µM and 5.16 µM TMSA52 two hours prior to challenge. Vehicle treated mice and mice treated with 5.16 µM TMSA52 lost an equivalent amount of weight by 3 dpi, while mice receiving 25.8 µM two hours prior to infection, or 258 µM at 2 or 24 hours prior to infection were wholly protected from morbidity (Figure 4B). A cohort of mice was sacrificed at 4 dpi to quantify viral burden in the lung, and compared to vehicle treated mice, those receiving 5.16 µM or 25.8 µM TMSA52 had a two- and three-fold decrease in viral burden, respectively (Figure 4C). In contrast, mice receiving 258 µM TMSA52 24 hours before challenge showed an identical decrease in viral burden to mice receiving the 258 µM TMSA52 2 hours before challenge, with 3/5 mice having undetectable viral loads by 4 dpi (Figure 4C). Together, these data illustrate that TMSA52 can persist in the airways to sustain protection against SARS-CoV-2 for at least 24 hours prior to challenge and can provide comparable protection against weight loss and viral burden at a 10-fold lower dose when administered 2 hours prior to infection.

**Fig. 4.**
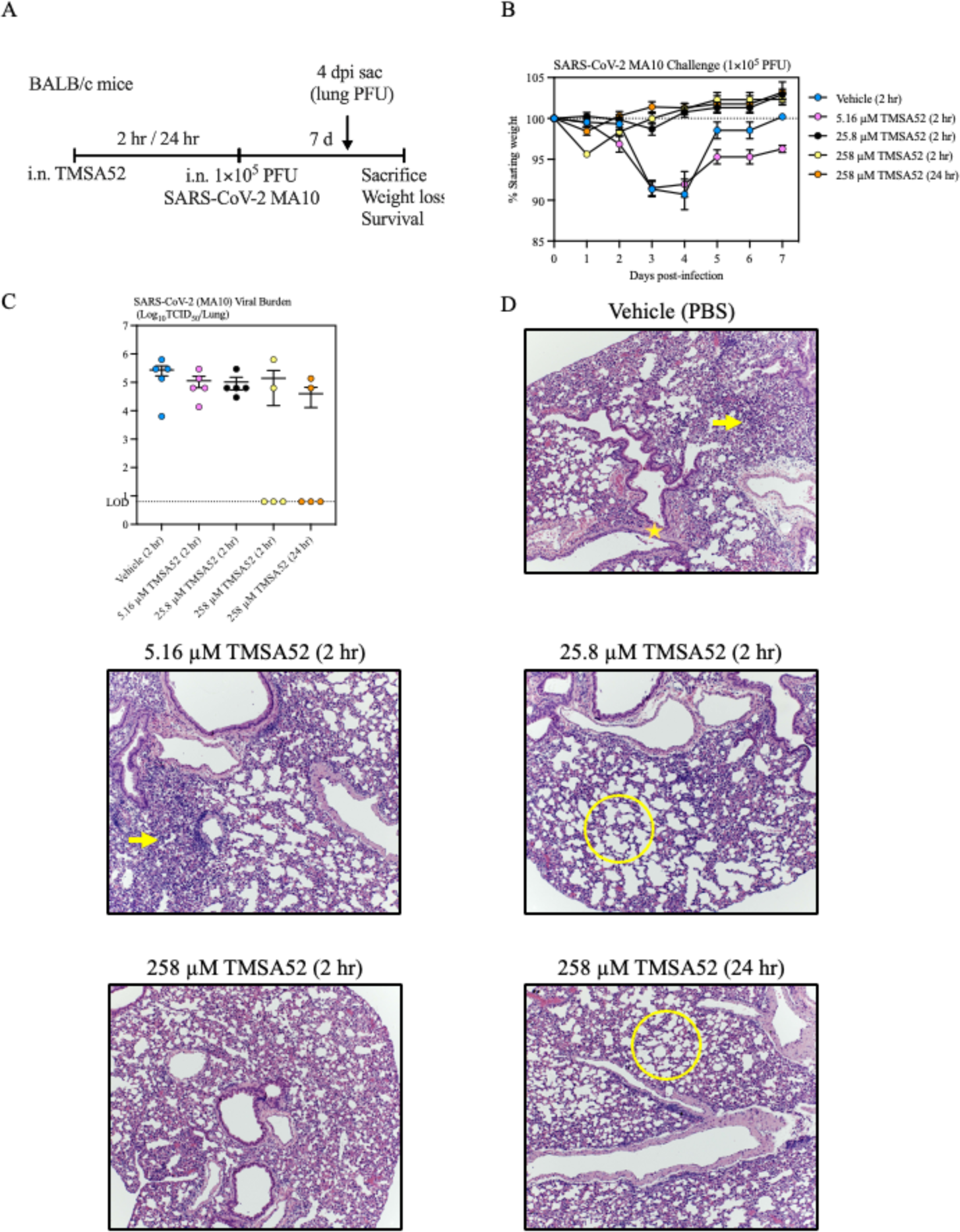
Protective assessment of different doses of intranasally delivered aptamers against challenge with SARS-CoV-2. (**A**). Animals were treated intranasally (i.n.) with either nuclease-free PBS (vehicle), or TMSA52 either 2 or 24 hours prior to challenge with 1×10_5_ PFU SARS-CoV-2 MA10. Animals were subsequently monitored for weight loss and mortality for seven days with a sub cohort sacrificed four days post-infection for lung viral titers and histopathology. (**B**) Weight loss following SARS-CoV-2 MA10 challenge. (**C**) Lung viral burden of a cohort of animals (n = 5) sacrificed four days post-infection. (**D**) Hematoxylin and eosin-stained lung sections from a cohort of animals sacrificed four days post-infection with yellow arrows indicating areas of diffuse epithelial damage, yellow stars representing epithelial sloughing, and yellow circles representing healthy lung architecture. Data is presented as mean ± SEM. Data is representative of one independent experiment, n = 5 mice/group.

We went on to perform H&E staining on lung slices from infected animals at 4 dpi. The lungs of mice treated with vehicle and 5.16 µM TMSA52 showed prominent epithelial damage and sloughing (yellow arrows and star, respectively; Figure 4D, top right, middle left). Mice treated with 25.8 µM TMSA52 two hours prior to infection, and those receiving 258 µM 24 hours before infection had few signs of pathology, in agreement with the absence of weight loss in those groups. (Figure 4D, middle right, bottom right). As expected, mice administered 258 µM TMSA52 two hours prior to infection had largely preserved lung architecture (yellow circles) and showed minimal pulmonary damage compared to vehicle, and 5.16 µM TMSA52 mice in congruence with the absence of substantial weight loss (Figure 4D, bottom left). Collectively, these data demonstrate that TMSA52 can protect mice against morbidity and mortality following SARS-CoV-2 infection at lower doses, and at protracted time points.

### Respiratory mucosal therapy with aptamer nanomaterials protects against SARS-CoV-2

Currently authorized mAbs for SARS-CoV-2 are administered intravenously to individuals with symptomatic disease and at high risk of progressing to severe COVID-19 (*5*). Having demonstrated that prophylactic respiratory mucosal administration of aptamer nanomaterials provides robust protection against SARS-CoV-2 infection equivalent to a monoclonal antibody therapy, we next sought to determine if they provide commensurate protection when administered therapeutically post-infection. To this end, BALB/c mice were infected with a SARS-CoV-2 MA10 and 24 hours post-infection were i.n. administered TMSA52 or IrNP@TMSA52. A subset of animals was treated intraperitoneally (i.p.) with S309 to recapitulate its systemic delivery in humans (Figure 5A). Consistent with previous results (Figure 2B, C), vehicle treated mice rapidly lost body weight and 1/5 mice reached endpoint by 4 dpi (Figure 5B, C). Unlike previous experiments, mice treated with S309 experienced pronounced weight loss, but all mice began to recover by 4 dpi (Figure 5B). Mice treated with TMSA52 showed no protection against weight loss, and all mice succumbed to infection by 4 dpi (Figure 5B, C). In contrast, animals that received IrNP@TMSA52 began to recover lost weight by 3 dpi, one day earlier than S309 treated mice, outperforming the archetype mAb therapy (Figure 5B, C). In agreement with the substantial weight loss and mortality observed, TMSA52 treated mice showed no decrease in viral load at 4 dpi (Figure 5D). However, IrNP@TMSA52 treatment reduced pulmonary viral burden 7.7-fold relative to vehicle treated mice, while S309 only decreased viral titers 2.7-fold relative to vehicle treated mice (Figure 5D). Notably, while marked epithelial damage (yellow arrows) was observed in the lungs of vehicle, TMSA52, and S309 treated animals (Figure 5E), IrNP@TMSA52 treated mice displayed reduced lung pathology and improved pulmonary architecture (yellow circles, Figure 5E, bottom left). The above data indicate that topical administration of IrNP@TMSA52 after infection protected against severe disease and lung pathology relative to standard-of-care parenteral administration of monoclonal antibodies. This highlights the potential of aptamer nanomaterials not just as a prophylactic agent, but as a potential therapeutic to mitigate clinical disease, reduce viral burden, and pathology when administered as a therapeutic.

**Fig. 5.**
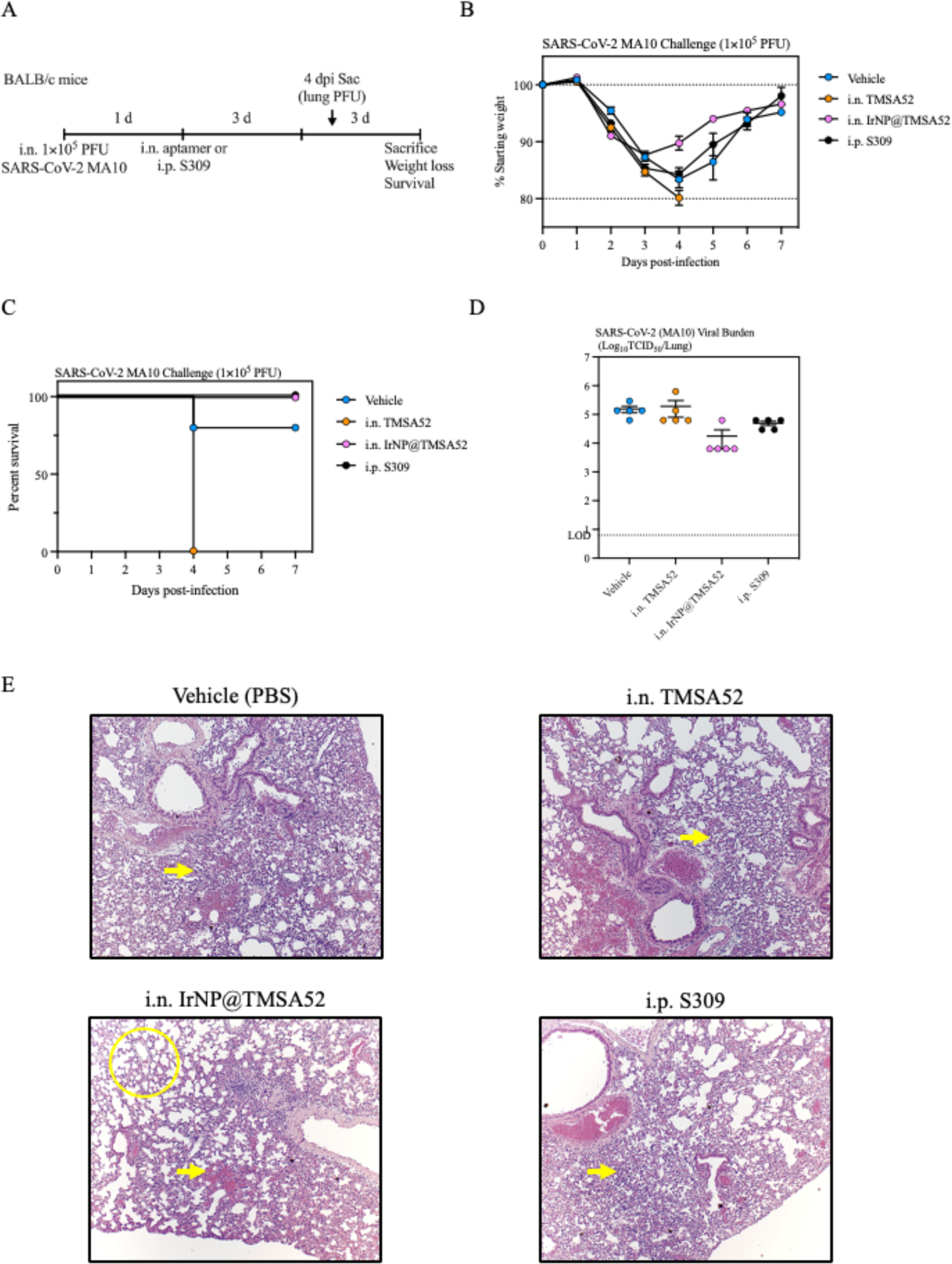
Protective assessment of therapeutic intranasally delivered aptamers. (**A**). Animals were challenged with 1 × 10_5_ PFU of SARS-CoV-2 MA10 and 24 hours post-infection mice were treated intranasally (i.n.) with either nuclease free PBS (vehicle), TMSA52, IrNP@TMSA52, or intraperitoneally (i.p.) with S309. Animals were subsequently monitored for weight loss and mortality for seven days with a sub cohort sacrificed four days post-infection for lung viral titers and histopathology. (**B**) Weight loss following SARS-CoV-2 MA10 challenge. (**C**) Survival following SARS-CoV-2 MA10 challenge. (**D**) Lung viral burden of a cohort of animals (n = 5) sacrificed four days post-infection. (**E**) Hematoxylin and eosin-stained lung sections from a cohort of animals sacrificed four days post-infection with yellow arrows indicating areas of diffuse epithelial damage, and yellow circles representing healthy lung architecture. Data is presented as mean ± SEM. Data is representative of one independent experiment, n = 5 mice/group.

### Respiratory mucosal prophylaxis with aptamer nanomaterials protects against a contemporary Omicron sub-variant

The continued emergence of increasingly disparate SARS-CoV-2 Omicron sub-variants has rendered many mAb therapies including Sotrovimab ineffective. TMSA52 has been well-characterized to bind a wide-array of SARS-CoV-2 variant spike proteins *in vitro*, however its *in vivo* utility against SARS-CoV-2 VoC had not been tested (*15*). To test whether TMSA52 could mediate protection against antigenically distant SARS-CoV-2 VoC, k18-hACE2 mice were treated i.n. with nuclease-free PBS (vehicle), TMSA52, or S309 two hours prior to infection with a lethal dose of SARS-CoV-2 XBB.1.5 (Figure 6A) (*31*). Vehicle treated mice rapidly lost weight, and all mice progressed to endpoint within 5 dpi (Figure 6B, C). Mice treated with S309 were protected from weight loss and mortality despite abrogated neutralizing capacity against XBB.1.5, likely due to enrichment of S309 in the airways after i.n. compared to the typical i.v. administration in clinical settings. Though 3/5 TMSA52 treated mice lost some weight following infection, all mice recovered to their original starting weight by 9 dpi and survived (Figure 6B, C). Vehicle treated mice had consistently high lung viral burden at 4 dpi. In contrast with earlier observations upon ancestral SARS-CoV-2 challenge, S309 and TMSA52 failed to significantly reduce pulmonary viral burden at 4 dpi, though there was a trend towards reduced titers in 4/5 TMSA52-treated mice (Figure 6D).

**Fig. 6.**
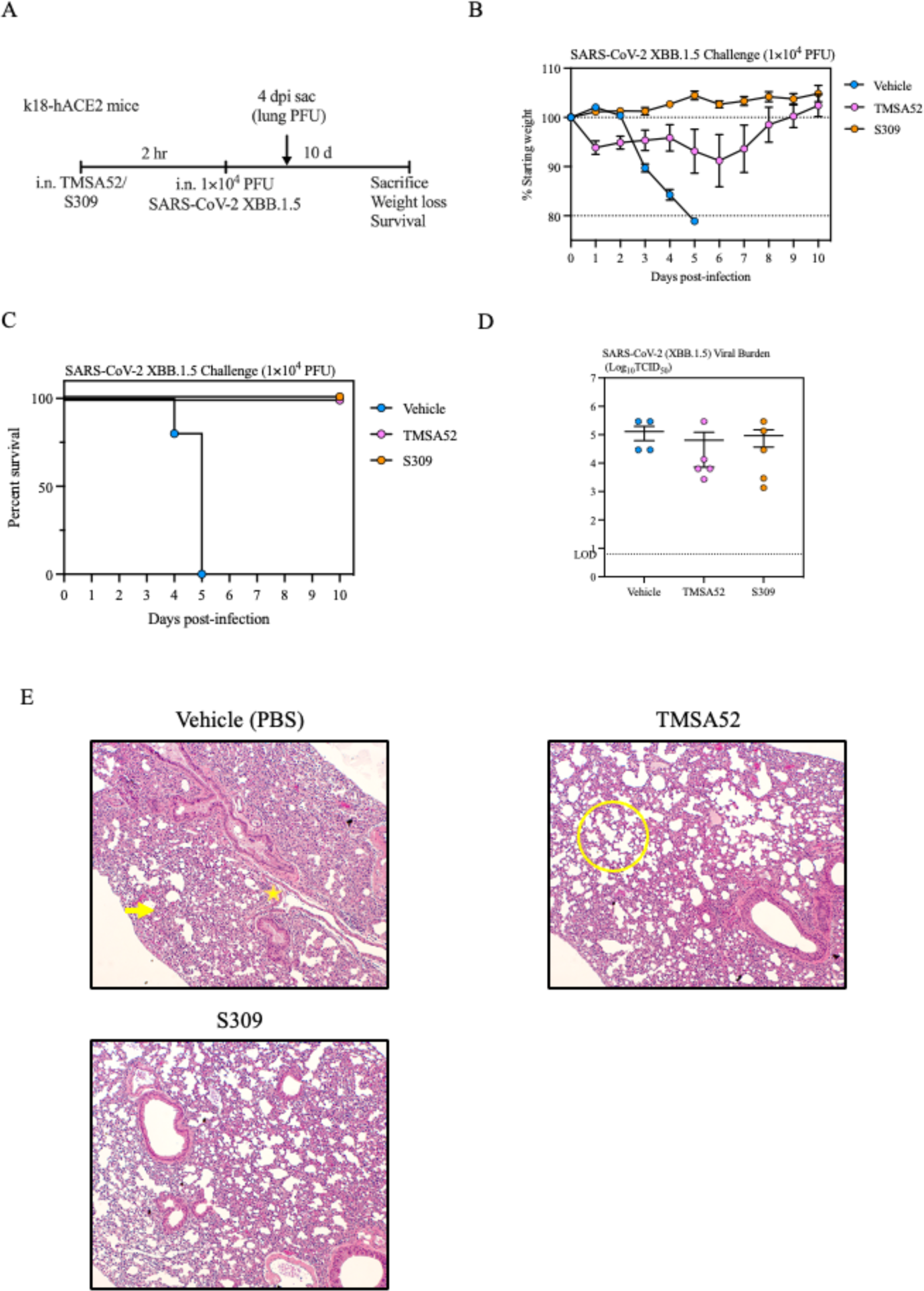
Protective assessment of intranasally delivered aptamers against lethal challenge with SARS-CoV-2 Omicron XBB.1.5. Animals were treated intranasally (i.n.) with either nuclease free PBS (vehicle), TMSA52, or S309 two hours prior to challenge with a lethal dose (1 × 10_4_ PFU) of SARS-CoV-2 XBB.1.5. Animals were subsequently monitored for weight loss and mortality for 10 days with a sub cohort sacrificed four days post-infection for lung viral titers and histopathology. (**B**) Weight loss following SARS-CoV-2 XBB.1.5 challenge. (**C**) Survival following SARS-CoV-2 XBB.1.5 challenge. (**D**) Lung viral burden of a cohort of animals (n = 5) sacrificed four days post-infection. (**E**) Hematoxylin and eosin-stained lung sections from a cohort of animals sacrificed four days post-infection with yellow arrows indicating areas of diffuse epithelial damage, yellow stars representing epithelial sloughing, and yellow circles representing healthy lung architecture. Data is presented as mean ± SEM. Data is representative of one independent experiment, n = 5 mice/group.

H&E staining on lung slices of vehicle treated mice at 4 dpi showed pronounced epithelial damage (yellow arrow) and epithelial sloughing (yellow star; Figure 6E, top left). In stark contrast, lungs from TMSA52-treated animals had markedly improved lung architecture comparable to S309 treated animals (Figure 6E, top right, bottom left). Collectively, the above data suggest that respiratory mucosal administration of aptamer nanomaterials can provide robust protection against even disparate SARS-CoV-2 VoC.

## DISCUSSION

While vaccines are often considered the best option for prophylaxis against infectious diseases, there is a major gap in prophylactic options for high-risk individuals who respond poorly to vaccines on the basis of being immunocompromised (e.g., transplant recipients, those on certain chemotherapies, frail elderly individuals, etc.), or for whom vaccines are contraindicated. Monoclonal antibodies have emerged as a key frontline countermeasure against viral pathogens. However, as was the case for monoclonal antibodies that received emergency use approval during COVID-19, they can rapidly lose efficacy due to antigenic drift (*32*). In addition to waning efficacy against variant viruses, traditional mAb therapies are further constrained by high discovery and manufacturing costs, maintenance of cold-chain, and a high degree of clinical expertise and infrastructure to support i.v. administration (*7, 8*). This underscores the necessity for novel therapeutics that maintain effectiveness despite viral evolution and are amenable to less burdensome routes of administration.

While DNA aptamers have been widely lauded for their value as diagnostic sensors, their utility as anti-infective agents for viral infections have thus far been limited to *in vitro* applications due to poor stability and pharmacokinetic properties for common methods of delivery (i.e., oral, intravenous). However, these limitations can be overcome by alternative modes of delivery. Indeed, both clinically approved therapeutic aptamers (Macugen and Izervay) are administered by intraocular injection. To our knowledge, we are the first to describe respiratory mucosal administration of a DNA aptamer as an agent to protect against SARS-CoV-2 *in vivo*. Respiratory mucosal administration of TMSA52 and IrNP@TMSA52 prevented weight loss and mortality following lethal infection with SARS-CoV-2 with comparable efficacy to a clinically-approved monoclonal antibody. In addition to preventing clinical disease, both aptamer nanomaterials prevented lung pathology, and reduced pulmonary viral burden, with IrNP@TMSA52 providing sterilizing protection in the lung.

In this study, TMSA52, a trimeric DNA aptamer previously identified for its broad-spectrum recognition of antigenically diverse SARS-CoV-2 spike proteins, was evaluated as a versatile and universal anti-SARS-CoV-2 countermeasure. TMSA52 readily binds the spike proteins across all major SARS-CoV-2 VoCs, including Omicron. The trimeric structure of TMSA52 is designed to mirror the trimeric spike protein, significantly enhancing its binding efficiency over monovalent MSA52 (*15*). Aptamers have undergone clinical evaluation for a variety of disease indications, and two modified RNA aptamers that target human proteins have been approved by the FDA. The first, Macugen, binds vascular endothelial growth factor to treat age-related macular degeneration (AMD), while Izervay recognizes complement C5, and is a treatment for geographic atrophy secondary to AMD (*33, 34*). Notably both are administered intravitreally to their site of action, likely due to the ephemeral half-life of aptamers in serum (*29*). DNA aptamers offer several possible advantages as anti-viral countermeasures: the SELEX process is a fast and efficient way to rapidly identify aptamers for any new target, they are more economical to produce than monoclonal antibodies, production is rapidly scalable, and aptamers offer remarkable stability under ambient conditions (*10*).

Moreover, the ability to rapidly tune the valency of aptamers provides immense flexibility and the opportunity to enhance effectiveness by increasing avidity. Conjugation of MSA52 onto the phosphoramidite trebler increased aptamer stability, binding affinity, and resilience against antigenic changes in the spike protein. Likewise, conjugation of TMSA52 to nanoparticles (iridium nanoplates, or spherical gold nanoparticles) further decreased the IC_50_ in neutralizing assays. Interestingly, both aptamer nanomaterials were more resilient to antigenic changes in spike than S309, the parental antibody of Sotrovimab, and maintained neutralization against antigenically divergent variants such as Omicron BA.5, and XBB.1.5.

A favorable safety profile is crucial for any inhaled therapeutic to ensure it does not interfere pulmonary function. While mAbs have a lengthy history of being well tolerated for clinical use, aptamer-based biologics have only thus far been approved for intravitreal administration in humans. In our study, we have shown that respiratory mucosal administration of TMSA52 did not induce pathogenic responses in the airways before or after SARS-CoV-2 infection. These findings collectively indicate that respiratory mucosal delivery of TMSA52 was safe and well tolerated.

In the context of infectious disease countermeasures, dose sparing can be an important way to maximize availability of treatments when case numbers are high. We therefore performed a dose de-escalation study and found comparable protection against morbidity in mice given a 10-fold lower dose (25.8 µM) than that at which we began our studies (258 µM). This highlights another advantage of respiratory mucosal administration of biologics, as local delivery provides high concentrations of the product directly at the site of infection (*35*). Juxtaposed to this, i.v. infusion of mAb therapeutics requires high doses to be administered, as concentrations of antibodies within the lung are up to 500-fold lower than in the serum (*36*). In addition to dose finding, it is important to determine the duration of efficacy provided by a potential prophylactic. Thus, we extended the interval between aptamer administration and infection, and found equivalent protection against both morbidity and viral burden for TMSA52 when administered up to one day in advance of virus challenge. Further investigation is needed to determine if the therapeutic window extends beyond 1 day prior to infection, and if protection at these protracted time points is preserved with lower doses of aptamer (e.g. 25.8 µM). It also remains unknown what dose would be relevant in a human lung, that has a surface area of 85 m^2^ in comparison to the murine lung which is several orders of magnitude less at 82.2 cm^2^ (*37, 38*). In addition, it remains to be delineated how preservation processes such as spray-drying might affect aptamer stability for long term storage at ambient conditions, and how these particles may be distributed throughout the RM via inhaled aerosol or the use of a dry powder inhaler.

Unexpectedly, TMSA52 administered therapeutically (post-infection) contributed to intensified weight loss relative to vehicle treated mice, and increased mortality in the absence of heightened viral titers or lung pathology. The mechanism behind this remains unclear, and further airway immune phenotyping following therapeutic administration of aptamer nanomaterials post-infection is required. In contrast, therapeutic delivery of IrNP@TMSA52 offered improved protection relative to systemically delivered S309, with mice recovering one day earlier and exhibiting less lung pathology than S309 treated mice.

Finally, we show that respiratory mucosal TMSA52 retained its protective efficacy in highly susceptible k18-hACE2 mice infected with the antigenically distant and contemporary SARS-CoV-2 XBB.1.5 variant. Though TMSA52 treated mice lost some weight following infection, all mice recovered to their original starting weight and survived. Moreover, both TMSA52 and S309 partially decreased viral burden and lung pathology post-infection. Together, these findings suggest that aptamer-based nanomaterials represent a new class of anti-viral countermeasures.

Another unique possibility for aptamer-based platforms is the potential use the same aptamer as both a diagnostic and therapeutic modality. Current lateral-flow antigen tests for SARS-CoV-2 mainly recognize the nucleocapsid protein from a nasopharyngeal swab (*39*). However, this has no direct predictive value for the efficacy of treatments that may be administered upon diagnosis. In contrast, if TMSA52 can recognize the SARS-CoV-2 spike protein in a rapid antigen test (at least at some empirically determined threshold), then it infers that TMSA52-therapeutics should also be capable of offering protection.

In summary, we have developed next-generation aptamer nanomaterials (TMSA52 and IrNP@TMSA52), that neutralize a diverse range of SARS-CoV-2 antigenic variants with little fluctuation in potency. We demonstrate that these aptamer nanomaterials can be delivered to the respiratory mucosa and are capable of similar protection against infection as that mediated by monoclonal antibodies. This provides proof-of-concept for aptamers as a viable new therapeutic platform for protection against respiratory virus infections. The cost and manufacturing benefits of this platform warrant its exploration as a core component of future pandemic preparedness arsenals.

## MATERIALS AND METHODS

### Study design

The objective of our study was to establish the neutralization properties of aptamer-based nanomaterials as novel anti-infective agents against SARS-CoV-2 in mice. We describe a novel one-pot synthesis to affix trimeric DNA aptamers onto iridium nanoplates, and test their neutralizing potency against SARS-CoV-2 VoC. We next evaluated aptamer-based nanomaterials as a prophylactic agent delivered i.n. to the respiratory mucosa of mice (n = 5 per treatment group), to prevent weight loss, mortality, pathology, and decrease viral burden. Animals were pooled then randomized prior to treatment. Humane endpoint was determined to be a loss of >20 % of their starting body weight. Lung viral burden was quantified by TCID_50_ using the Spearman-Kärber method. Evaluation of lung pathology was performed in a blinded manner. We investigated the immunostimulatory properties of the aptamer nanomaterials by quantifying the inflammatory cytokine milieu via multiplex, and described the innate immune cells recruited to the airway by flow cytometry. We sought to determine an effective dose, and the durability of protection offered by the aptamer nanomaterials when administered i.n. prior to infection. In addition, we assessed the aptamer nanomaterials as a therapeutic agent delivered following SARS-CoV-2 infection. Finally, we appraised the potential of prophylactic i.n. aptamer administration prior to challenge with a divergent SARS-CoV-2 VoC.

### Chemicals and reagents

DNA oligonucleotides listed in Table S1 were obtained from Yale University (New Haven, CT, United States), McMaster University (Hamilton, Canada), or Integrated DNA Technologies (Coralville, IA, United States) and purified using 10 % denaturing polyacrylamide gel electrophoresis (dPAGE) containing 8 M urea. Sodium borohydride (NaBH_4_, 98 %), sodium hexachloroiridate (III) hydrate (Na_3_IrCl_6_ · H_2_O, M.W. = 473.9), potassium phosphate monobasic (KH_2_PO_4_, ≥ 99 %), sodium phosphate dibasic (Na_2_HPO_4_, ≥ 99 %), potassium chloride (KCl, ≥ 99 %), sodium chloride (NaCl, ≥ 99.5 %), 4-(2-hydroxyethyl)-1-piperazineethanesulfonic acid (HEPES, ≥ 99 %), magnesium chloride (MgCl_2_, ≥ 99 %), acetic acid (HOAc, ≥ 99.7 %), 3,3‘,5,5‘-tetramethylbenzidine (TMB, > 99 %), sodium acetate (NaOAc, ≥ 99%), sulfuric acid (H_2_SO_4_, 95– 98 %), hydrogen peroxide solution (30 % H_2_O_2_), dimethylformamide (DMF), sodium hydroxide (NaOH, ≥ 98 %), hydrochloric acid (HCl, 37 %), streptavidin (Cat. No. SA101), bovine serum albumin (BSA, Cat. No. A7906), and tween-20 were all obtained from Sigma–Aldrich (St. Louis, MO, United States). The pseudotyped lentiviruses expressing the spike proteins of ancestral (Cat. No. 79981), B.1.1.7 (Cat. No. 78158), B.1.351 (Cat. No. 78160), P.1 (Cat. No. 78159), B.1.617.2 (Cat. No. 78216), B.1.1.529 (Cat. No. 78349), BA.2.12.1 (Cat. No. 78646), and BA.5 (Cat. No. 78652) SARS-CoV-2 were purchased from BPS Bioscience (San Diego, CA, United States). Dulbecco’s modified Eagle’s medium (DMEM, Gibco), fetal bovine serum (FBS, Gibco), trypsin-EDTA (Gibco), penicillin-streptomycin (Gibco), and Hoechst 33342 (Cat. No. 62249) were acquired from ThermoFisher Scientific (Ottawa, Canada). 96-well microtiter plate (clear and black, polystyrene, flat bottom) was from Celltreat Inc (Pepperell, MA, United States). Ultrapure water (Milli-Q System, Millipore-Sigma, Etobicoke, Canada) was used to prepare all aqueous solutions.

### Synthesis of IrNPs

Briefly, freshly prepared Na_3_IrCl_6_ (100 µL, 10 mM) and NaBH_4_ (100 µL, 100 mM) aqueous solution were sequentially dispensed into ultrapure water (1 mL) in a 1.5 mL centrifuge tube. The tube was sealed immediately, followed by vigorous vortexing for 10 seconds. The mixture was then shaken at 200 rpm at room temperature (22°C) for 10 hours. IrNPs were obtained after centrifugation at 3000 RCF (relative centrifugal force) for 5 minutes and washed with water three times. To scale up IrNP synthesis, Na_3_IrCl_6_ (4 mL, 10 mM) and NaBH_4_ (4 mL, 100 mM) aqueous solution were added to ultrapure water (40 mL) in a 50 mL centrifuge tube, followed by procedures described above to obtain IrNPs. After three washes with ultrapure water, IrNPs were dried under vacuum, followed by measuring weight with an analytical balance. IrNP weight was measured to be 4.492 mg, which corresponds to 93.6 µg/mL IrNPs in 48 mL solution. According to equation, m_Ir_ = ρ_Ir_ × v_Ir_, where m_Ir_ represents the mass of each IrNP, ρ_Ir_ denotes the density of iridium, which is 22.56 g/cm^3^, and v_Ir_ is the volume of each IrNP, calculated to be 1.495 × 10^-13^ cm^3^ (based on the side length of IrNPs: 0.985µm and the thickness of IrNPs: 59.3nm), m_Ir_ is determined to be 3.373 × 10^-12^ g. According to equation c = m / (m_Ir_ × NA × v), where c signifies the concentration of IrNP, m indicates the total mass of IrNPs (4.492 mg), NA is Avogadro’s number 6.02 × 10^23^ mol^-1^, v represents the total volume of IrNPs (48 mL), and c is calculated to be 46.1 fM.

### Characterization of IrNPs

The SEM analyses were conducted on Hitachi S-4700 FE-SEM operating at 20 kV. The TEM, SAED patterns, and EDX analyses were performed on a JEOL microscope (JEOL 2010F) at 200 kV. HRTEM images were obtained on an FEI Titan 80-300 microscope operating at 300 kV. XPS spectra were acquired from an SSX-100 system (Surface Science Laboratories, Inc.). XRD measurement was conducted on a Scintag XDS2000 powder diffractometer. UV-Vis absorbance of samples in the microtiter plate was obtained by a plate reader (Tecan, Switzerland).

### Preparation of IrNP@TMSA52

IrNPs (1 mL, 46.1 fM) and streptavidin (10 µL, 16.7 µM) were mixed in PBS and incubated at 22°C for 1 hour, followed by incubation at 4°C for 24 hours. After washing twice with PBS, biotinylated trimeric aptamer TMSA52-B (100 µL, 3 µM) in PBS was added and incubated at 22°C for 1 hour. The aptamer conjugated IrNPs (IrNP@TMSA52) were washed twice with PBS and stored at 4°C before use. The preparation of IrNP-MSA52 followed the same procedure as that of IrNP@TMSA52, with the only difference being the replacement of TMSA52-B with MSA52-B. IrNP@TMSA52 was analyzed with a 2 % agarose gel. Briefly, monomeric aptamer MSA52, trimeric aptamer TMSA52, and IrNP@TMSA52 with the same amount of aptamer (10 µL, 100 nM) were loaded on 2 % agarose gel containing 1 × fluorescent dye SYBR Safe DNA Gel Stain (Invitrogen), run at 150 V for 45 minutes, and analyzed by Amersham Typhoon Biomolecular Imager (ThermoFisher Scientific, Waltham, MA, United States). To determine the number of TMSA52-B conjugated on each IrNP, a FAM-labelled complementary DNA (10 µL, 20 µM, FAM-AS listed in Table S1) was used to hybridize with IrNP@TMSA52 (100 µL, 46.1 fM) in PBS for 30 minutes. After three washes with PBS, the IrNP conjugates were resuspended in PBS containing 0.05 % tween-20 and 8 M urea, followed by denaturation at 90°C for 10 minutes. After centrifugation, the supernatant was collected to measure the fluorescence intensity. The concentration of TMSA52-B conjugated on IrNPs was calculated by comparing the fluorescence intensity to a standard curve to determine the number of TMSA52-B on each IrNP. The number of MSA52-B conjugated on each IrNP was also determined using the same procedure.

### Detection of pseudoviruses using IrNP@TMSA52

To assess the universal recognition ability of IrNP@TMSA52 for SASR-CoV-2 variants, IrNP@TMSA52 was utilized for the detection of different pseudoviruses. Pseudoviruses were detected by IrNP@TMSA52-based enzyme-linked aptamer binding assay (ELABA). IrNP@TMSA52 was first prepared as previously described. After washing once with PBS, IrNP@TMSA52 were blocked with 10 % BSA in PBST (PBS containing 0.05 % tween-20) by incubation at 37°C for 1 hour and then 4°C for 24 hours before use. Subsequently, a streptavidin-coated 96-well microtiter plate was prepared by addition of streptavidin (100 µL, 5 µg/mL) in PBS to the wells, followed by incubation at 4°C for 12 hours. Biotinylated trimeric aptamer TMSA52-B (100 µL, 200 nM) in PBS was then conjugated with streptavidin by incubation at 22°C for 30 minutes. After two washes with PBS, the plate was blocked with BSA (100 µL, 2 %) in PBS by incubation at 37°C for 1 hour. Different concentrations of pseudoviruses (100 µL) in sample dilution buffer (SDB, PBST with 0.1 % BSA) were added and incubated at 22°C for 1 hour. The plate was washed twice with SDB. Whereafter, IrNP@TMSA52 (100 µL, 46.1 fM) in SDB was introduced and incubated at 22°C for 1 hour, followed by four washes with PBST. Finally, substrate solution (100 µL, 0.8 mM TMB, 2 M H_2_O_2_, 0.1 M HAcO/NaAcO buffer, pH = 4) was added to the wells and reacted at 22°C for 20 minutes. The catalytic reaction was terminated with H_2_SO_4_ (20 μL, 2 M). The absorbance of oxidized product at 450 nm was measured using a plate reader (Tecan, Switzerland).

### Investigation of aptamer nanomaterial stability in culture medium

IrNP@TMSA52 (10 µL, containing 1000 nM aptamer) was mixed with 90 % Dulbecco’s Modified Eagle Medium (DMEM, containing 10 % FBS, 90 µL), followed by incubation at 37°C for different times. After denaturing at 90°C for 10 minutes, the aptamers were analyzed by electrophoresis using 5 % PAGE gel with 8 M urea in 1 × TBE at 150 V for 45 minutes. The stability of monomeric aptamer MSA52-B and trimeric aptamer TMSA52-B in DMEM was also investigated as described for IrNP@TMSA52.

### Neutralization of pseudoviruses by IrNP@TMSA52

HEK 293T cells (100 µL, 5000 cells) expressing ACE2 were cultured with DMEM in a 96-well plate for 12 hours. Different concentrations of IrNP@TMSA52 (10 µL) in DMEM were incubated with pseudoviruses (10 µL) at 37°C for 1 hour. The IrNP@TMSA52 and pseudovirus mixture was then added to 293T cell monolayer. After incubation for 6 hours, the cell culture medium was refreshed, followed by cell culture at 37°C for 48 hours for GFP expression. Hoechst (10 µL, 20 µM) was added to stain cell nuclei by incubation at 37°C for 20 minutes. The cells were washed once with PBS (100 µL) before fluorescence imaging analysis.

### Cell lines

Vero E6 (CRL-1586, American Type Culture Collection (ATCC), Manassas, VA, United States) were cultured at 37°C in Dulbecco’s Modified Eagle medium (DMEM, McMaster University, Hamilton) with 10 % fetal bovine serum (FBS, Gibco, Massachusetts, United States), 1 % HEPES pH = 7.3 (McMaster University, Hamilton), 1 % Glutamax (Invitrogen, Massachusetts, United States) and 100 U/mL of penicillin–streptomycin (Invitrogen, Massachusetts, United States).

### *In vitro n*eutralization assay with SARS-CoV-2

Vero E6 cells were seeded at a density of 2.5 × 10^4^ cells/well in clear flat-bottom TC-treated 96-well plates (Falcon, 353377, Tewksbury, MA, United States) with DMEM fortified with 10 % FBS, 1 % Penicillin-Streptomycin, 1 % HEPES (pH = 7.3), 1 % Glutamax. and incubated at 37°C, 5 % CO_2_ overnight., Aptamer constructs were serially diluted starting with a concentration of 75 µM of MSA52, 25 µM of TMSA52 and 10 µM of IrNP@TMSA52 for Ancestral (SB3), and the XBB.1.5 Omicron variant. These aptamer nanomaterial dilutions were incubated with SARS-CoV-2 (330 PFU/well) for 1 hour at 37°C, 5 % CO_2_. Following this incubation, the mixture was transferred onto Vero E6 cells and re-incubated for 1 hour under the same conditions. The mixture was replaced with identical dilutions of the aptamers and incubated for 24 hours at 37°C, 5 % CO_2_. Post-incubation, cells were fixed with 3.7 % paraformaldehyde and incubated at 4°C. Plates were blocked for 1 hour at 37°C with reagent diluent (0.5 % bovine serum albumin (BSA), 0.02 % sodium azide, in 1× Tris-Tween buffer). Following a 1 hour incubation at 37°C, plates were washed three times with 1× Tris-Tween wash buffer. After washing, rabbit anti-SARS/SARS-CoV-2 Nucleocapsid Polyclonal Antibody (Invitrogen, Massachusetts, United States) was diluted 1:1000 in reagent diluent. Plates were incubated for 1 hour, at 37°C, followed by three washes with 1× Tris-Tween buffer. A goat anti-rabbit IgG (H+L), biotinylated secondary antibody (Invitrogen, Massachusetts, United States) diluted 1:8000 in reagent diluent was added for 1 hour at 37°C. Plates were subsequently washed three times prior to the addition of streptavidin-alkaline phosphatase (1:2000, Southern Biotech, Birmingham, AL, United States) to all wells for 1 hour at 37°C. After washing, pNPP one component microwell substrate solution (Kementec, Denmark) was added to each well and plates were developed for 10 minutes. The reaction was quenched with an equal volume 3 N sodium hydroxide. The optical density (O.D.) at 405 nm was read on a Biotek Synergy H1 (ThermoFisher Scientific Waltham, MA, United States). O.D. values were normalized based uninfected control wells and percent neutralization was calculated based on the ratio between sample O.D. value to infected control value.

### SARS-CoV-2 viruses

SARS-CoV-2 strain SB3 was provided by Dr. Arinjay Banerjee, Dr. Karen Mossman, Dr. Samira Mubareka, and Dr. Rob Kozak (Banerjee et al., 2020). SARS-CoV-2 strain MA10 was generously provided by Dr. Ralph Baric (Leist et al., 2020). SARS-CoV-2 strain hCoV-19/USA/MD-HP40900/2022 (Lineage XBB.1.5, NR-59104, Dr. AS Pekosz) were obtained from BEI Resources (Manassas, VA, United States).

### Aptamer/antibody treatment and infection

Animals were anesthetized with isoflurane and intranasally administered 40 µL of TMSA52, IrNP@TMSA52, or S309 monoclonal antibody diluted in nuclease-free PBS. Mice were infected with 1 × 10^4^ or 1 × 10^5^ PFU SARS-CoV-2 administered intranasally in a final volume of 40 µL diluted in PBS. For intraperitoneal administration of S309, mice were administered 1.5 mg/kg S309 diluted in nuclease-free PBS to a final volume of 100 µL per mouse. Mice were monitored for clinical signs and weight loss daily, with 80 % of initial weight considered humane endpoint, in accordance with institutional guidelines.

### SARS-CoV-2 viral burden determination in tissues

Lung and brains were homogenized using a Bead Mill 24 homogenizer (ThermoFisher Scientific Waltham, MA, United States). Homogenates were clarified by centrifugation at 300 × g and frozen at -80°C. Homogenates were thawed, and serially diluted 1:10 in serum-free DMEM supplemented with 1% HEPES pH = 7.3, 1 mM sodium pyruvate, 1 % L-Glutamine and 100 U/mL of penicillin– streptomycin. 100 mL of viral inoculum was plated on Vero E6 cells in 96-well plates (4 × 10^4^ cells/well) for 1 hour at 37°C, 5 % CO_2_, at which point the inoculum was replaced with low-serum DMEM supplemented with 2 % FBS, 1 % HEPES pH = 7.3, 1 mM sodium pyruvate, 1 % L-Glutamine and 100 U/mL of penicillin–streptomycin. Wells were assessed for cytopathic effect at 5 dpi using an EVOS M5000 microscope (ThermoFisher Scientific Waltham, MA, United States).

### Cytokine analysis

BAL samples were concentrated through PierceTM Protein Concentrators with a 50 kDa molecular weight cut-off (MWCO) (ThermoFisher Scientific Waltham, MA, United States) according to manufacturer’s instructions, with volumes normalized prior to concentration. Evaluation of BAL cytokines were performed by Eve Technologies (Calgary, Canada) through a mouse cytokine array/chemokine array 44-plex (MD44).

### Statistical analysis

A one-way ANOVA with Tukey’s multiple comparisons was used to test the significant differences for *in vitro* neutralization against SARS-CoV-2, and to test the differences in pulmonary viral burden following murine challenge. All data points are presented as mean ± SEM.

## LIST OF SUPPLEMENTARY MATERIALS

Supplemental fig. 1. Neutralization of pseudoviruses expressing the SARS-CoV-2 BA.5 spike by aptamers and aptamer-conjugated nanomaterials.

Supplemental fig. 2. Synthesis and characterization of hexagonal IrNPs.

Supplemental fig. 3. Characterization of IrNPs.

Supplemental fig. 4. Characterization and analysis of IrNP@TMSA52.

Supplemental fig. 5. Flow cytometry gating strategy.

Table S1. DNA oligonucleotide sequences from this manuscript.

## Acknowledgements

Authors are thankful to Dr. Ralph Baric for provision of mouse-adapted SARS-CoV-2 (MA10). The following reagent was obtained through BEI Resources, NIAID, NIH: SARS-CoV-2, strain hCoV-19/USA/MD-HP40900/2022 (Lineage XBB.1.5), NR-59104. Authors are grateful to the McMaster Histology Core Facility, the McMaster Flow Cytometry Core Facility, the McMaster Central Animal Facility, the McMaster Canadian Centre for Electron Microscopy Facility, and the McMaster Biosafety Office.

This work was funded by an Alliance Grant from Natural Sciences and Engineering Research Council of Canada (NSERC) to YL (grant number ALLRP 570428-2021) and a research grant from Canadian Institutes of Health Research (CIHR) to Y.L. (grant number GA5-17777777)

M.S.M. was funded, in part, by a Canada Research Chair in Viral Pandemics and research funding from Zentek Ltd.

## Author contributions

MRD, JL, SA, ZZ, JG, AM, JA, YL, and MSM conceived and designed the study. MRD, JL, ZZ, JG, AM, JA, and SA performed experiments and analyzed data. MRD, JL, SA, YL, and MSM wrote the paper.

## Competing interests

Provisional patents have been filed with the United States Patent and Trademark Office for the use of trimeric aptamers as diagnostic and therapeutic agents. The patent applications are owned by McMaster University and under license to Zentek through a license agreement that covers diagnostic, neutralization and therapeutic use of trimeric aptamers.

## Data and materials availability

All data, code, and materials described within are available upon request from the corresponding authors.

---

**Supplemental fig. 1.**
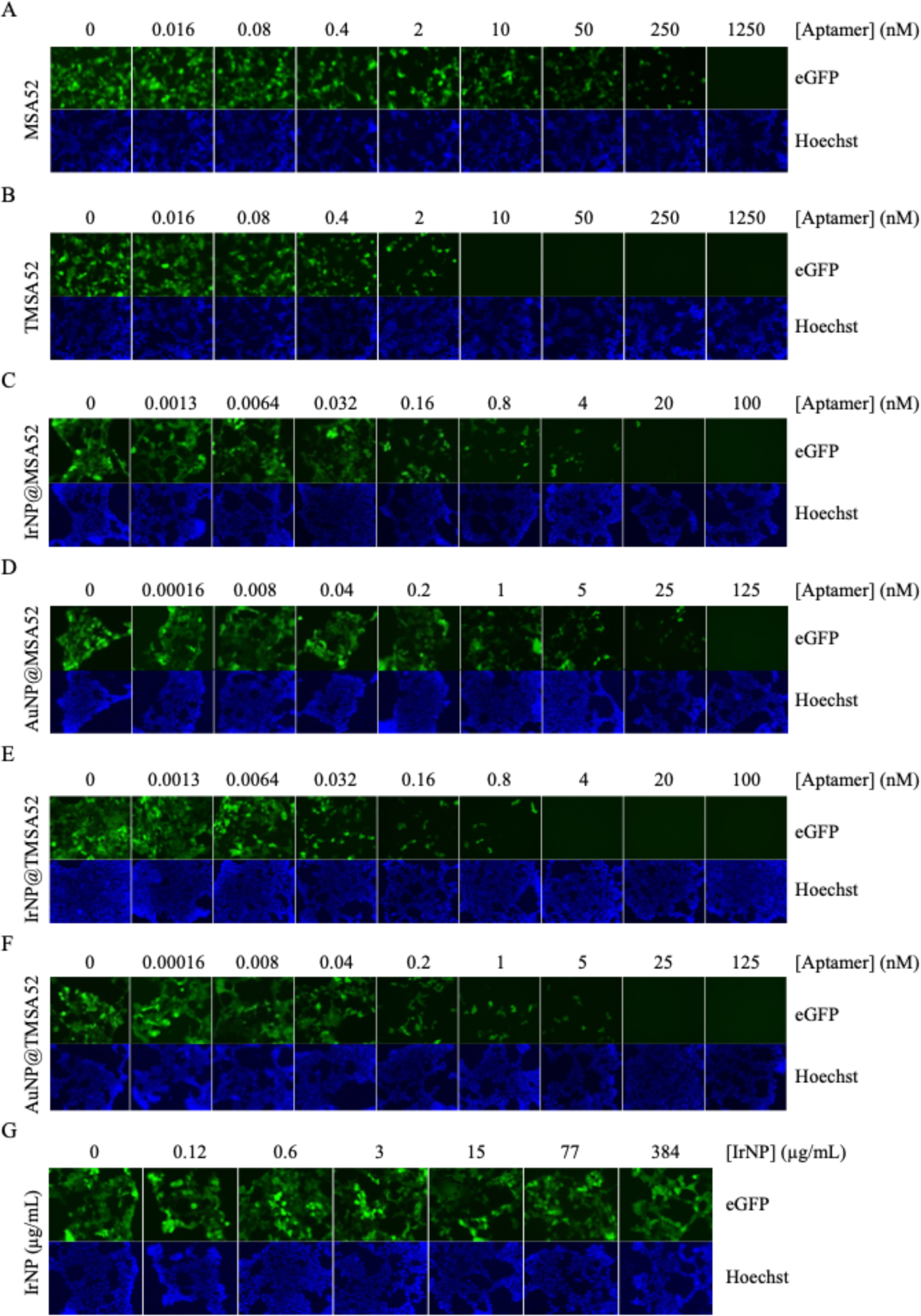
Neutralization of pseudoviruses expressing the SARS-CoV-2 BA.5 spike by aptamers and aptamer-conjugated nanomaterials. (**A**) Fluorescence images showing the infection of HEK 293T cells by SARS-CoV-2 BA.5 spike expressing pseudoviruses in the presence of monomeric aptamer MSA52, (**B**) trimeric aptamer TMSA52, (**C**) IrNP@MSA52, (**D**) AuNP@MSA52, (**E**) IrNP@TMSA52 (E), (**F**) AuNP@TMSA52, or (**G**) IrNP alone.

**Supplemental fig. 2.**
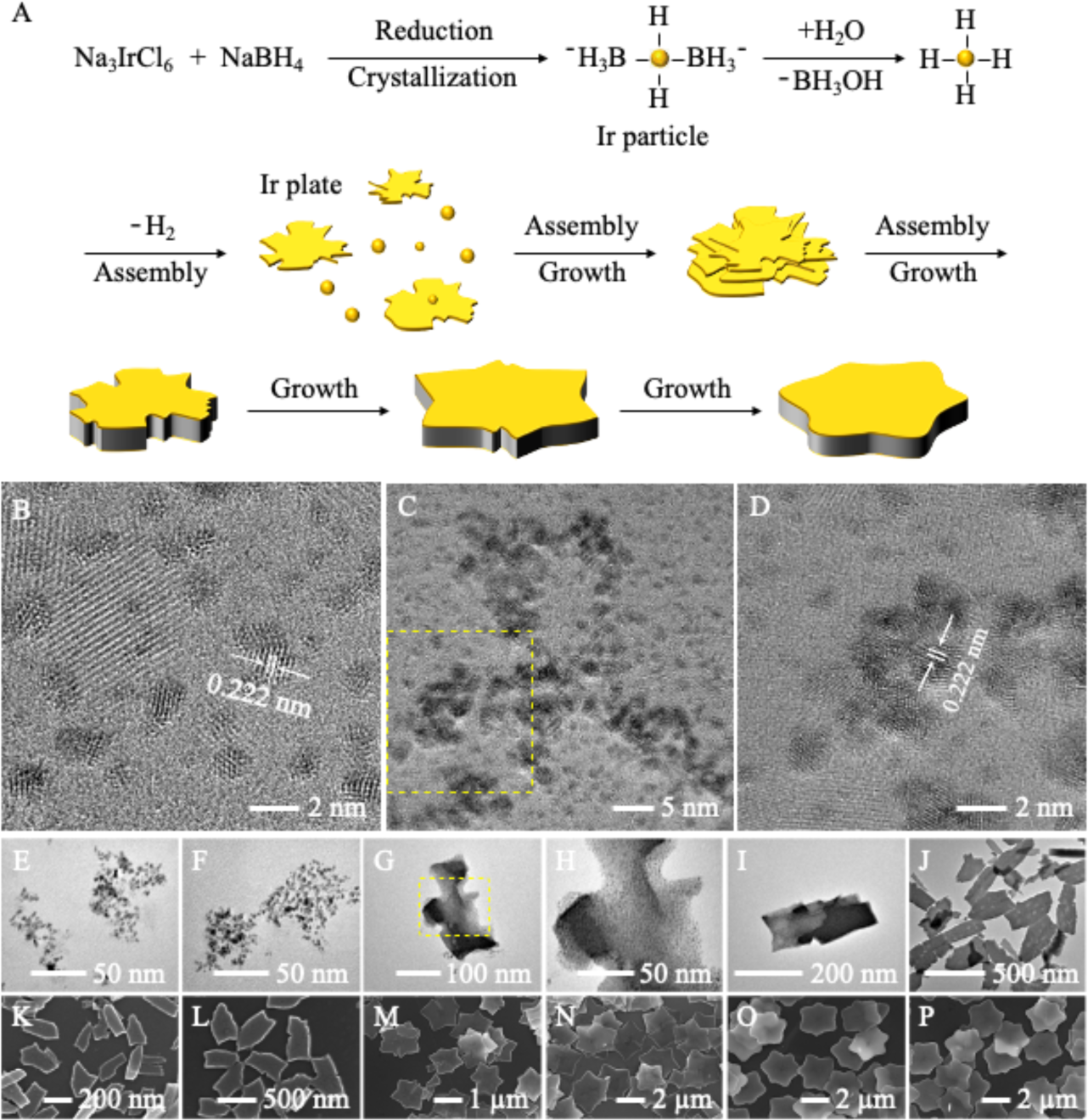
Synthesis and characterization of hexagonal IrNPs. (**A**) Na_3_IrCl_6_ is first reduced by NaBH_4_ to form Ir nanoclusters, which is stabilized by borohydride anions through electronic repulsion. Due to the excellent catalytic activity of Ir towards the hydrolysis of borohydride anions and the generation of H_2_, Ir nanoclusters assemble into Ir nanosheets that furthers the formation of multilayer assembly. Irregular IrNPs are then obtained via an assembly/growth mechanism, which finally transform into hexagonal IrNPs. (**B**) Ir nanoclusters before the formation of IrNPs were observed with a particle size of 1.24 ± 0.34 nm and 0.222 nm lattice fringes, corresponding to the {111} planes of fcc Ir. (**C, D**) Numerous crystal boundaries are readily discernible within the initial 2-minute timeframe of the synthesis process, providing clear evidence of the assembly of Ir nanoclusters into Ir nanosheets. (**E, F**) The single-layer Ir nanosheets grow gradually extending their sizes during 5 to 10 min. (**G, H**) After 20 minutes, the presence of multilayered Ir nanosheets becomes evident as Ir nanoclusters consistently assemble onto the lateral surfaces of the Ir nanosheets. (**I, L**) The assembly/growth processes proceed progressively with multilayer Ir nanosheets transforming into irregular nanoplates. (**M, N**) Hexagonal nanoplates with radial wedge-shaped gaps. (**O, P**) Hexagonal nanoplates.

**Supplemental fig. 3.**
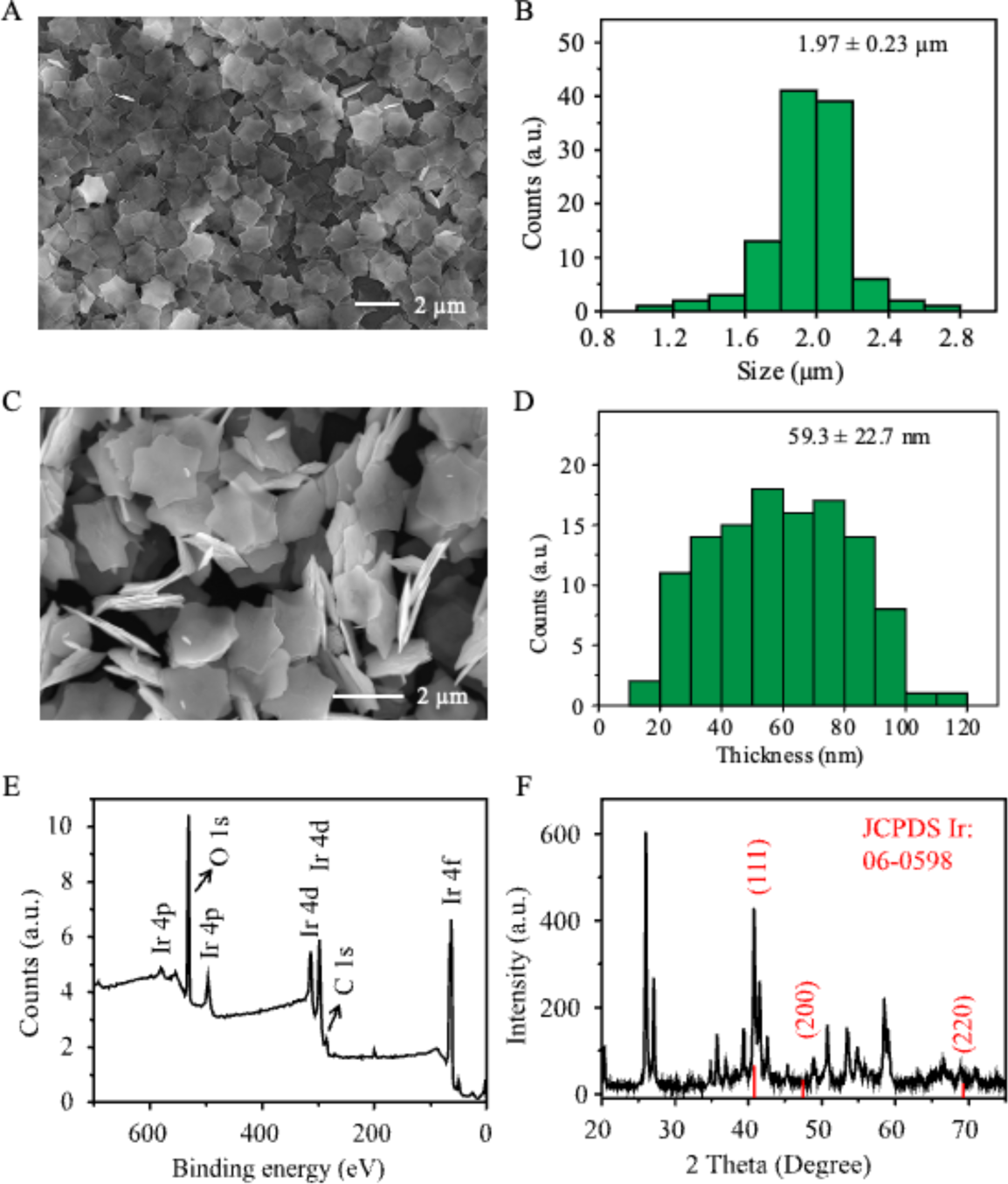
Characterization of IrNPs. (**A**) Upon synthesis completion, free-standing uniform hexagonal IrNPs were obtained with a typical yield above 95%. (**B**) Statistical analyses showed that IrNPs had a size distribution of 1.97 ± 0.23 µm and a thickness of 59.3 ± 22.7 nm (**C, D**). (**E**) The elemental composition of IrNPs was characterized by X-ray photoelectron spectroscopy (XPS) exhibiting that IrNPs are dominated by Ir. (**F**) Many narrow diffraction peaks were observed in X-ray powder diffraction (XRD) analysis, which is attributed to the crystal structure of IrNPs with large size. Compared with typical fcc Ir materials (JCPDS card No. 06-0598), a peak located at 40.6°is ascribed to the {111} planes of fcc Ir. The disappearance of two peaks derived from {200} and {220} planes of fcc Ir and observation of several unknown peaks are possibly due to the directional assembly of Ir nanoclusters during synthesis and multilayered superlattice structures of IrNPs.

**Supplemental fig. 4.**
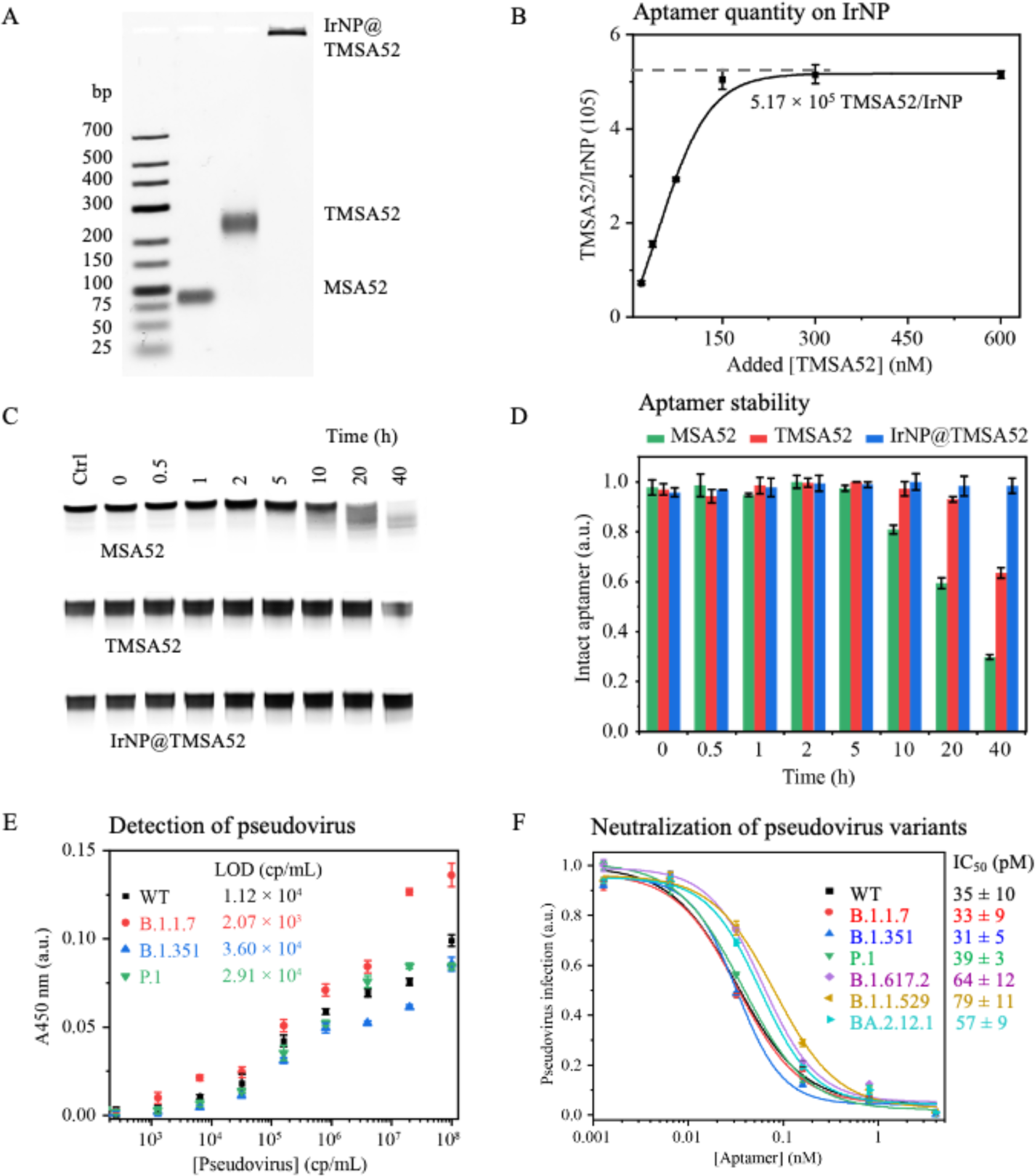
Characterization and analysis of IrNP@TMSA52. (**A**) Agarose gel analysis of monomeric aptamer (MSA52), trimeric aptamer (TMSA52), and TMSA52 affixed to iridium nanoplate (IrNP@TMSA52). (**B**) Determination of TMSA52 quantity conjugated on each IrNP. (**C**) dPAGE and (**D**) corresponding plots showing the stability of monomeric aptamer MSA52, trimeric aptamer TMSA52, and IrNP@TMSA52 in Dulbecco’s Modified Eagle Medium (DMEM) supplemented with 10% fetal bovine serum (FBS) at 37°C. (**E**) Plots depicting the results of the IrNP@TMSA52-based ELABA (Enzyme-Linked Aptamer Binding Assay) for detecting pseudoviruses expressing spike proteins from different SARS-CoV-2 variants, including wild-type (WT), B.1.1.7, B.1.351, and P.1. (**F**) Determination of IC_50_ values of IrNP@TMSA52 for the neutralization of pseudoviruses expressing the spike protein from different SARS-CoV-2 variants of concern.

**Supplemental fig. 5.**
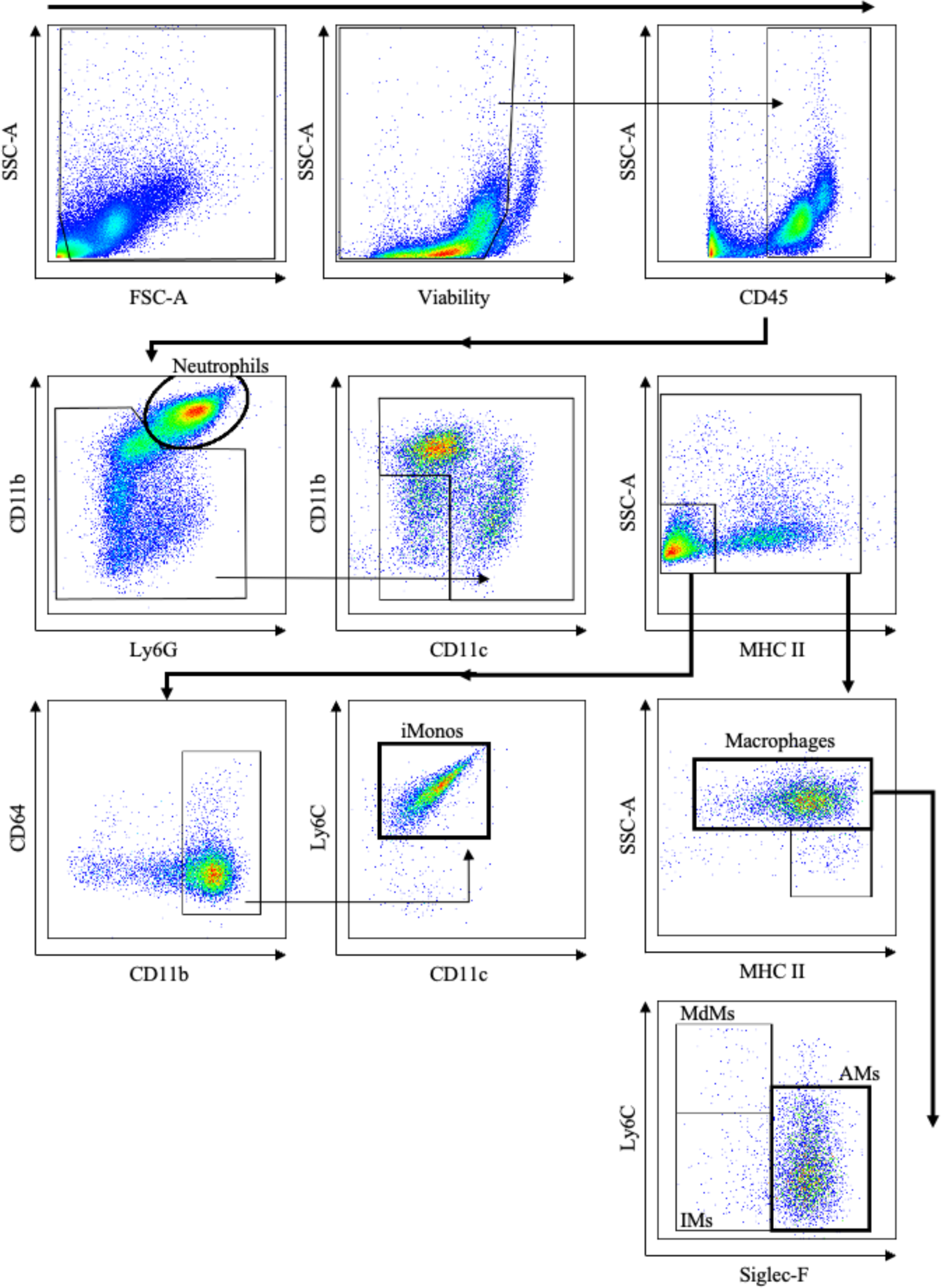
Flow cytometry gating strategy. Gating strategy applied in this study to differentiate various innate immune cell populations including neutrophils, inflammatory monocytes (iMonos), monocyte-derived macrophages (MdMs), interstitial macrophages (IMs), and alveolar macrophages (AMs) in the airways via a comprehensive flow cytometry panel.

**Table S1.**
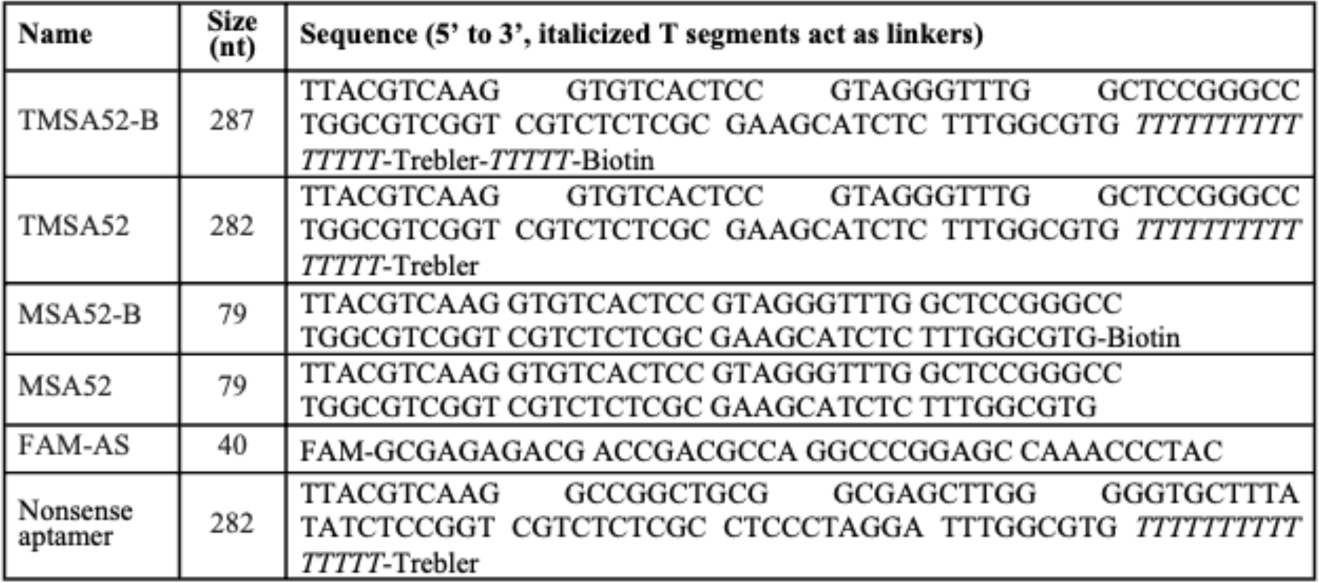
DNA oligonucleotide sequences from this manuscript.

